# The Nonstructural Proteins 3 and 5 from Flavivirus Modulate Nuclear-Cytoplasmic Transport and Innate Immune Response Targeting Nuclear Proteins

**DOI:** 10.1101/375899

**Authors:** Margot Cervantes-Salazar, Ana L. Gutiérrez-Escolano, José M. Reyes-Ruiz, Rosa M. del Angel

**Affiliations:** Departamento de Infectómica y Patogénesis Molecular, CINVESTAV-IPN, México, D.F., México

## Abstract

Viruses hijack cellular proteins and components to be replicated in the host cell and to evade the immune response. Although flaviviruses have a cytoplasmic replicative cycle, some viral proteins such as the capsid (C) and the RNA dependent RNA polymerase, NS5, can reach the nucleus of the infected cells. Considering the important roles of NS5 in viral replication and in the control of the immune response, and its striking presence in the nucleus, the possible functions of this protein in some mechanisms orchestrated by the nucleus was analyzed. We isolated and identified nuclear proteins that interact with NS5; one of them, the DEAD-box RNA helicase DDX5 is relocated to the cytoplasm and degraded during infection with DENV, which correlates with its function in IFN dependent response. Since DDX5 and many other proteins are relocated from the nucleus to the cytoplasm during flavivirus infection, the integrity and function of the main regulator of the nuclear-cytoplasmic transport, the nuclear pore complex (NPC) was evaluated. We found that during DENV and ZIKV infection nucleoporins (NUPs) such as TPR, Nup153, Nup98, and Nup62 were cleavaged/degraded. The protease NS2B-NS3 induces NUPs degradation and it causes a dramatic inhibition of mature mRNAs export to the cytoplasm but not the export of DDX5 protein, which is dependent on NS5. Here we describe for the first time that the NS3 and NS5 proteins from flavivirus play novel functions hijacking the NPC and some nuclear proteins relevant in triggering immune response pathways, inducing a favorable environment for viral replication.

**IMPORTANCE:** Viruses, as intracellular obligate parasites, hijack cellular components to enter and replicate in infected cells. Remarkably, in many cases, viruses hijack molecules with crucial functions for the cells. Here it is described how RNA viruses such as DENV and ZIKV, with a cytoplasmic replicative cycle, use NS3 and NS5, two of their unique non-structural proteins with enzymatic activity, to modulate nuclear-cytoplasmic transport. We found that NS3 disrupts the nuclear pore complex, the main regulator in nuclear-cytoplasmic transport, causing a strong reduction in the amount of mature mRNAs in the cytoplasm and an inhibition in innate immune response. Additionally, NS5 induces the relocation of nuclear proteins to the cytoplasm such as DDX5, involved in immune response, which is later degraded by NS3. These findings allow the understanding of crucial mechanisms that viruses use to deal with the control of the immune response to grant the production of new viral particles.

## INTRODUCTION

In the last years, the world has experienced several outbreaks of different arbovirus such as dengue (DENV), chikungunya, Zika (ZIKV), West Nile (WNV), yellow fever (YFV), and Mayaro, among others. However, the most important viral disease transmitted by mosquitoes in the world is dengue. This infection is present in more than 100 countries, and each year more than 390 millions of dengue infections are reported^1^. DENV and ZIKV, members of the *Flaviviridae* family and *Flavivirus* genus, are single-stranded and positive polarity RNA virus. Its genomic RNA has a unique open reading frame that encodes for a polyprotein, which is cleaved by viral and cellular proteases in three structural proteins (capsid, C, prmembrane, prM and envelope, E) and seven nonstructural proteins (NS1, NS2A, NS2B, NS3, NS4A, NS4B, and NS5). Flavivirus as many other viruses are obligate intracellular parasites. Every step of the viral life cycle, from entry to viral release, requires close interaction with cellular proteins. Interestingly, the first interaction of the virus with host cell receptor is able to trigger a serial of dynamic processes that allow, viral internalization, translation, the formation of replicative complexes, replication and morphogenesis and at the same time control of immune response. Therefore, viruses hijack and manipulate cellular proteins, components, and processes to be replicated in the host cell. Consequently, a perfect balance between the efficient viral replication and the control of immune response grant a productive viral life cycle. Flaviviruses use different strategies to evade immune response; one of them is the inhibition of IFN response mediated by at least, in the case of DENV, of the nonstructural proteins NS2A, NS4A, NS4B, and NS5 which target the signal transducer and activator of transcription proteins 1 and 2^2^. A second mechanism to inhibit immune response is common for many RNA viruses and is the use of the nonstructural viral proteins to modify the endoplasmic reticulum (ER) membranes to generate partially isolated compartments known as replication complexes (RC), where the new viral particles are replicated and formed^2^. The RCs create a barrier that minimizes the detection of double-stranded RNA or 5’-phosphorylated RNA in the cytoplasm. Interestingly, the formation of the RC requires fatty acids and cholesterol. To this regard, the NS3 protein sequesters the fatty acids synthase to the RC and viral infection increases the activity of the HMG CoA reductase (limiting enzyme in the cholesterol synthesis pathway). The activation of the HMG CoA reductase is a consequence of the inhibition of the molecule considered as the cellular metabolism controller, the AMP-dependent kinase (AMPK)^3^.

Although there is an evident hijack of relevant cytoplasmic cell proteins, components and organelles during flaviviral infection, the hijack of nuclear components and the effect in cellular expression is poorly understood. Even though flaviviral infection takes place in the cytoplasm, two viral proteins are also located in the nuclei of infected cells; one is the C protein and the other, the NS5 protein. NS5 is the largest and most conserved viral protein (105 kDa and 900 amino acids)^4, 5^ which contains three domains: the RNA dependent RNA polymerase, the RNA guanylyltransferase, and the methyltransferase, and is required for the synthesis and capping of new viral genomic RNAs^6–9^. Additionally, NS5 can modulate the host immune response^10–13^ and cellular splicing^14^.

Given the importance of NS5 protein in DENV viral replication it could be predicted that the location of the protein was the ER, where viral replication takes place; however, this protein is also present in the nucleus of the infected cells^15^. In this regard, Pryor et al. described that NS5 contains a nuclear localization sequence in a lysine-rich domain, which is responsible for its import to the nucleus by the importin α/β (Imp α/β)^16^. Moreover, when mutations in the lysine-rich domain were included, a dramatic decrease in the production of infective virus particles was detected. In contrast, in a recent report, the C-terminal 18 amino acid region of DENV NS5 has been reported to be involved in the regulation of its subcellular localization^17^. Additionally, in this region, authors found a conserved arginine residue essential for infectious virus production^17^. Interestingly, the nuclear import blockage induced by the Imp α/β inhibitor, ivermectin, dramatically inhibits viral infection, suggesting that the presence of NS5 in the nucleus is required during viral replicative cycle^16^. Although the apparent importance of the presence of NS5 in the nucleus, serotype differences in its nuclear localization exist. While the NS5 protein from DENV-2 and -3 accumulates in the nucleus, the NS5 protein from DENV-1 and -4 is predominantly cytoplasmic^15^. The importance of NS5 subcellular distribution remains unsolved and could be related to an auxiliary function. To this respect, in a recent report, the interaction of NS5 with components of the U5 small nuclear ribonucleoprotein (snRNP) was confirmed. This interaction reduces the efficiency of pre-mRNA splicing and renders an advantageous cellular environment for DENV replication^14^.

Considering the important roles of NS5 during DENV replication and in the control of immune response, and its presence in the nucleus, the cellular compartment in which the transcription regulation takes place, we decided to analyze its participation in some of the mechanisms orchestrated by the nucleus. For this purpose, we isolated and identified nuclear cellular proteins that interact with NS5. Among others, the DEAD-box RNA helicase DDX5 that belongs to a family of proteins involved in translation, ribosomal biogenesis and transcription, and splicing regulation, was identified as NS5 binding proteins. This protein is relocated to the cytoplasm and degraded in DENV infected cells. Given the relocation of DDX5 and of many other nuclear proteins to the cytoplasm, the integrity and function of the nuclear pore complex (NPC), the main regulator of the nuclear-cytoplasmic transport, was evaluated during DENV and ZIKV. Our results show, for the first time that the NS3 and NS5 proteins play new undescribed functions inhibiting dramatically the mRNA export and interferon response by targeting nuclear proteins.

## MATERIALS AND METHODS

### Cell culture and virus propagation

Huh-7 cells, a differentiated hepatocyte-derived cellular carcinoma cell line, (provided by Dr. Ana Maria Rivas, Universidad Autónoma de Nuevo León, México), were grown in advanced DMEM supplemented with 2 mM glutamine, penicillin (5 x 10^4^ U/ml) streptomycin (50 μg/ml), 10% fetal bovine serum (FBS), 1 ml/L of amphotericin B (Fungizone) at 37°C and a 5% CO_2_ atmosphere. Propagation of DENV serotype 2 New Guinea strain, was carried out in CD1 suckling mice brains and titers were determined by focus assays in Huh-7 cells. Propagation of ZIKV was performed in Vero cells and titers were determined by focus forming unit assays in Huh-7 cells. CD1 suckling mice brains lysates from mock-infected mice were used as control.

### Focus forming assay

Huh-7 cells grown in culture plates (1X10 ^5^ cells/plate) were infected with serial dilutions of DENV or ZIKV, and the infection was permitted for 24 or 48 hrs. Cells were fixed with 2% formaldehyde for 20 min, treated with permeabilizing solution (1% serum, and 0.2% saponin in PBS) for 20 min at room temperature (RT) and incubated with a mouse anti-E (4G2) antibody overnight at 4°C. Cells were incubated with anti-mouse coupled to FITC IgG and focus were visualized by fluorescence microscopy. Viral titer was expressed as focus forming units.

### Preparation of nuclear protein extracts

Huh-7 cells grown in culture plates (8×10^6^ cells/plate) for 24 hrs at 37°C were washed twice with 2 ml of PBS and 1 ml of detaching buffer (40 mM TRIS HCl pH 7.5, 2 mM EDTA, 150 mM NaCl). Cells were harvested, centrifuged at 700 rpm for 10 min at 4°C and the pellet was washed twice with 1 ml of buffer A (10 mM HEPES pH 7.9, 1.5 mM MgCl_2_, 10 mM KCl) and resuspended in 1 volume (100 or 200 μl) of buffer A in the presence of protease inhibitor cocktail (EDTA, EGTA and DTT Roche). Cells were lysed in a homogenizer at 4°C. Nuclei and debris were removed by centrifugation at 10,000 rpm for 30 min at 4°C and the supernatant was the cytoplasmic cell extract. The pellet was washed twice with 1 ml of buffer C (10 mM TRIS pH 7.8, 5 mM MgCl_2_, 10 mM KCl, 0.3 mM EGTA, 0.5 mM DTT, 0.3 M sucrose, and 2 mM ZnCl_2_) and centrifuged at 2500 rpm for 3 min. The pellet was resuspended in 200 μl of buffer D (20 mM TRIS pH 7.8, 5 mM MgCl_2_, 320 mM KCl, 0.2 mM EGTA, 0.5 mM DTT y 2 mM ZnCl_2_) in the presence of protease inhibitor cocktail. After 30 min incubation on ice, the mixture was sonicated for 10 seconds and cell extract was treated with 300 units of micrococcal nuclease (per 100μL of nuclear extract) in the presence of 5 mM of CaCl_2_ at RT for 15 min. Debris was removed by centrifugation at 14,000 rpm for 30 min at 4°C and the supernatant was the nucleic extract. The nuclear extracts used for western blot assays were prepared by using a subcellular fractionation kit (Thermos scientific). The cytoplasmic and nuclear extracts were quantified by BCA assay.

### Affinity chromatography with NS5

A total of 2 mg/ml of nuclear cell extract in interaction buffer (50 mM NaH_2_PO_4_ pH 8.0, 0.5 M NaCl y 10 mM imidazole) was pre-absorbed by incubation with 200 μl of Ni-NTA resin (in the absence of NS5 recombinant protein) for 3 hrs in gentle shaking. Pre-absorbed extracts were recovered by centrifugation at 3000 rpm for 5 min at 4°C. Next, 500 μl of the resin coupled with recombinant NS5 protein or 500 μl of the control resin (described before), were incubated with pre-absorbed cell extracts diluted in 5 volumes of interaction buffer overnight at 4°C with gentle shaking. The resins were recovered by centrifugation at 3000 rpm for 5 min and were subjected to four washes with 5 volumes of washing buffer to purify proteins natively containing increasing concentrations of imidazole and NaCl (6 mM imidazole, and 250 mM of NaCl; 7.5 mM imidazole, and 250 mM NaCl: 7.5 mM of imidazole, and 300 mM NaCl; 7.5 mM of imidazole, and 350 mM NaCl). Finally, the proteins bound to the recombinant NS5 or to the control resin were eluted with 0.50 M and 1 M of NaCl. The eluted proteins were precipitated overnight with acetone at -20 °C. The pellets were re-suspended in 10 mM TRIS, pH 7.4 and the proteins were separated by SDS-PAGE for 20 min at 80 volts. The gel was washed 3 times for 5 min with milliQ water, and fixed with F solution (40% methanol and 10% acetic acid) for 30 min. Proteins were stained with 50 ml Coomassie blue (blue R-250, Bio-Rad) for 30 min at RT, the bands were cut and analyzed by MALDI-TOF (ms/ms) in the Protein Core Lab Facility at Columbia University in New York, NY.

### Immunoprecipitation assay

Huh-7 cells were infected with DENV 2 at an MOI of 3, and harvested 24 hrs post infection. A total of 2 mg/ml of nuclear extract were pre-absorbed with 50 μl of protein G agarose (Roche) for 4 hrs at 4°C. The lysate was recovered by centrifugation at 1000 rpm for 3 min. An anti-DDX5, anti-hnRNP F, and anti-rabbit IgG (control) antibodies (Abcam) were coupled to 100 μl of protein G agarose beads by crosslinking with DCG (disuccinimidyl glutarate, Thermo Scientific). The protein G agarose coupled with the different antibodies was blocked in 200 μl of blocking buffer (1 mg/mL BSA in PBS), for 60 min at 4°C. Then, beads were incubated with the pre-absorbed cell lysate. The interaction was permitted overnight at 4°C with gentle shaking. The immunocomplexes were recovered by centrifugation at 1000 rpm for 3 min, and the beads were washed five times with 400 μl of IP buffer with proteases inhibitor (50 mM TRIS pH 8.0, 1% NP40, 150 mM NaCl, 10 mM EDTA). Proteins were eluted in 1M NaCl and precipitated with acetone overnight at -20°C. The pellet was re-suspended in 50 mM of Tris pH 7.4, and proteins were separated by SDS-PAGE, transferred to a nitrocellulose membrane (Bio-Rad), and analyzed by western blot.

### Classification and protein network analysis

To investigate the biological process, subcellular localization and signaling pathway associated with each identified protein, information from Swiss-Prot/TrEMBL database and DAVID Bio-informatics Resources (http://david.abcc.ncifcrf.gov/) were used. The proteins were also searched through REACTOME pathway database (http://www.reactome.org).

STRING network analysis of protein-protein interactions was performed to identify functionally linked proteins. The network is presented under confidence view, whereby stronger associations are represented by thicker lines or edges and vice versa, whereas proteins are represented as nodes.

### Western blot assay

Huh-7 cells were infected with DENV 2, DENV 4, or ZIKV, at an MOI of 3 and harvested at 12, 24, and 48 hours post infection (hpi). Cells were prepared in lysis buffer in the presence of protease inhibitor cocktail (ROCHE). A total of 50 μg/ml of total extract or immunoprecipitated proteins which had been heated or not for 10 min at 95°C in the presence of 3% β-mercaptoethanol, were separated by electrophoresis in 7.5% SDS-PAGE and transferred to nitrocellulose membrane (Bio-Rad). Membranes were blocked with 10% of non-fat milk in 0.5% PBS-Triton X-100. Cell proteins were detected by using an anti-DDX5 goat polyclonal antibody (1:5000, AbCam) or anti-hnRNP F rabbit polyclonal antibody (1:7000, AbCam) or anti-Nup62 rabbit polyclonal antibody (1:6000, AbCam) and anti-Nup98 rabbit polyclonal antibody (1:6000, Cell signaling) or anti-Nup153 mouse polyclonal antibody (1:3000, AbCam) or anti-TPR mouse monoclonal antibody (1:500, Santa Cruz Biotechnology, Santa Cruz, CA). The DENV viral proteins (NS3, NS5) or ZIKV (NS3) were detected by using rabbit polyclonal antibodies (1:5000 and 1:5000, GeneTex) respectively. An anti-rabbit HRP, anti-mouse HRP and anti-goat HRP antibodies (1:10000, Cell Signaling) were used as secondary antibodies in all cases. The proteins were visualized with Super Signal West Femto Chemiluminescent Substrate (Thermo scientific).

### Transfection assay

Huh-7 cells grown on slides were transfected with 3 μg of NS3-WT or NS3-S135A or NS5-HA plasmids from DENV2 (kindly donated by Dra. Ana Fernandez-Sesma and Dr. Adolfo García-Sastre) or with NS2B-NS3 plasmid from ZIKV (Kindly donated by Dr. Bowl) with Lipofectamine LTX (Invitrogen) according to the indications of the reagent, and 24 or 48 hrs post-transfection, cells were fixed for immunofluorescence assay.

### Immunofluorescences analysis

Huh-7 cells grown on slides were transfected or not with NS5-HA or NS3-HA or S135A-HA or NS3-ZIKV; the cells were infected or not with DENV2 or DENV4 or ZIKV at an MOI of 3. Twenty four hrs post transfection, the cells were treated or not with 30 μg/mL of Poly I:C (Polyinosinic:polycytidylic acid) for 24 hours and fixed at different hrs post infection with 3% formaldehyde (FA) for 20 min at RT. Cells were treated with permeabilizing solution (serum 1%, saponin 2mg/mL in PBS) for 20 min at RT. Cells were incubated with 1 μg/ml of either rabbit anti-NS5 protein, rabbit anti-NS3 protein, or mouse anti-E protein (4G2) antibodies and with antibodies directed against the cellular proteins Mab414 (AbCam), Nup62, Nup98, DDX5 or hnRNP F in permeabilizing solution, overnight at 4°C. Cells were incubated with 1 μg/ml of AlexaFluor 488-conjugated donkey anti-mouse IgG, AlexaFluor 555-conjugated goat anti-rabbit IgG, AlexaFluor 555-conjugated mouse anti-goat IgG or AlexaFluor 488-conjugated anti-rabbit IgG. Nuclei were stained with Hoechst (Santa Cruz Biotechnology, Santa Cruz, CA). Slides were observed in a Zeiss LSM700 laser confocal microscope.

### Immunoelectron microscopy analysis

Mock-infected and DENV-infected Huh7 cells were fixed with 4% paraformaldehyde/0.5% glutaraldehyde for 1 h at RT. Then, cells were dehydrated through increasing concentrations of ethanol, embedded in the acrylic resin (LR White) and polymerized under UV irradiation overnight at 4°C. The Resin-embedded cells sections of 70 nm were obtained and mounted on Formvar-covered nickel grids, incubated in PBS with 10% fetal bovine serum for 1 h to block nonspecific binding and reacted with anti-NS5 and anti-DDX5 antibodies diluted in 5% fetal bovine serum. The samples were washed three times and incubated with anti-rabbit or anti-goat IgGs antibodies conjugated to 15-nm (Ted Pella Inc., Redding, CA, USA) and 25-nm (Electron Microscopy Sciences) colloidal gold particles, respectively. Finally, sections were contrasted with uranyl acetate and lead citrate before being examined under a Jeol JEM-1011 transmission electron microscope. The mock-infected cells were treated under the same conditions as infected cells.

### FISH of poly (A)-containing RNAs

To detect the subcellular localization of the polyadenylated mRNAs, fluorescence in situ hybridization (FISH) assay was performed. Transfected or untransfected, infected or uninfected cells were fixed with 4% formaldehyde for 20 minutes. The formaldehyde was aspirated and cold methanol was added to each well for 10 min. Later, methanol was aspirated and 70% ethanol was added for 10 min. Finally, ethanol was aspirated and 1M Tris pH 8.0 was added for 5 min. After blocking the cells with 0.5% BSA for 20 min, the Cy5-Oligo-dT probe diluted in hybridization buffer at a final concentration of 1ng / mL (1mg/mL yeast tRNA, 0.5mM EDTA, 0.5% BSA, 10% Dextran sulfate and 25% formamide deionized) was added and incubated for 12 hrs at 37°C. Cells were washed 5 times with 4X SSC buffer and 5 times with 2X SSC buffer. The rabbit anti-NS3 antibody (1 μg /ml) was diluted in 2X SSC with 0.1% Triton X-100 and incubated overnight at 4°C. Cells were washed 5 times with 2X SSC and incubated with 1 μg/ml of AlexaFluor 555-conjugated goat anti-rabbit IgG. Nuclei were stained with Hoechst (Santa Cruz Biotechnology, Santa Cruz, CA). Slides were observed in a Zeiss LSM700 laser confocal microscope.

### *In silico* analysis

To predict the putative cleavage site by serine proteases or nuclear localization sequences (NLS) different software were used. To predict the putative cleavage site in Nup153, Nup62, hnRNP F and DDX5 (UniProtKB accession numbers: P49790 and P37198 or P52597 and P17844) and Nup98 (GenBank accession number: AAC50366.1) scan prosite database was used. We used the pattern, which represents two or three residues of Lys and/or Arg [KR] followed by any Ser, Gly or Ala^18^. The docking of NS2B-NS3 with Nup98 was analyzed by automatic docking ClusPro 2.0 Server and AutoDock vina. Docking was performed using 3D structure of Protein Data Bank (PDB) [PDB ID 2FOM and 2Q5X]^18^. The model was analyzed and visualized using the PyMOL program.

Nuclear Localization sequences (NLS) in NS3 protein from DENV and ZIKV were predicted by NLS-Mapper. A score of 8 was selected to identify the NLSs^19^. We used the Seq-NLS software from the University of South Carolina^20^ and LocNes software to analyze Nuclear Export Sequences^21^. The GenBank accession numbers of the sequences analyzed were: DENV-1 (AFN54943.1) DENV-2 (AAK67712.1), DENV-3 (AFN80339.1), DENV-4 (AEX09561.1) and ZIKV (YP_009227202.1).

### Treatment with protease inhibitors and cell viability assay

Huh-7 cells grown in 2.3 x 10^5^ cells/plate were infected at MOI of 3 for 8 hrs. Cells were incubated with two different serine protease inhibitors (Leupeptin 1 μM or TLCK.HCl 150μM) for 24 hrs. Cells were lysed and the integrity of the NUPs was analyzed by western blot assay using 60 μg of protein. Cells were lysed or fixed and the integrity of hnRNP F and DDX5, proteins was evaluated by western blot assay and confocal microscopy.

The viability of Huh-7 cells was evaluated by incorporation of propidium iodide in treated cells. The propidium iodide incorporation was measured by flow cytometry in a BDLCR-Fortessa X-20 apparatus and analyzed by Cyflogic version 1.2.1 software.

### siRNA mediated knockdown of DDX5 protein in Huh-7 cells

For siRNA-mediated knockdown of DDX5 expression, transfections were carried out according to the protocol recommended by the manufacturer (Santa Cruz Biotechnology, Santa Cruz, CA). Briefly, Huh-7 cells were plated in a 6-well plate to reach confluence; then, two mixtures were prepared: a) 50 or 100 nM of siRNA diluted in 150 μl of transfection medium (opti-MEM, Life Technologies) and b) 1.5 μl of siPORT (Applied Biosystems) in 100 μl of transfection medium and incubated 10 min at 37°C, combined by pipetting and incubated for 10 min. Next 250 μl of siRNA transfection medium was added to each tube and total volume was added to the cells. Cells were incubated in a CO_2_ incubator 8 hrs at 37°C, and the medium was aspirated and 200 μl of free medium containing 10% of fetal bovine serum was added. After 24 hrs post transfection, cells were transfected again under the same conditions and 24hrs after the second transfection, cells were treated or untreated with 30 μg/mL of Poly I:C (Polyinosinic:polycytidylic acid) for 24hrs. Finally, the cells were fixed and analyzed by confocal microscopy.

### Statistical analysis

Differences between the diverse treatments and control groups were evaluated using the statistical program Sigma-Plot 11 in all cases. One tail analysis of variance (ANOVA) was used. For all tests used a p≤0.05 was considered statistically significant.

## RESULTS

### Cellular proteins that interact with DENV2 NS5 in Huh7 cells

Many different strategies have been used to isolate and identify cellular proteins that interact with viral proteins, including double hybrid systems, immunoprecipitation and affinity chromatography assays among others. For the present analysis, we used an affinity chromatography assay with the recombinant DENV2 His tag-NS5 protein synthesized and purified (80% purity) from inclusion bodies by GenScript (New Jersey, USA). The protein was resuspended in buffer (20 mM PBS, 5% glycerol, 150 mM NaCl, pH 8.0) and quantified by Bradford protein assay (0.378 mg/ml). The recombinant NS5 protein was coupled to a NiNta column (Qiagen) and its presence and purity were confirmed in an SDS-PAGE gel stained with Coomassie blue (Figure 1A Supplementary material) and by western blotting using anti-NS5 and anti-His polyclonal antibodies (Figure 1B lanes 1 and 2 Supplementary material respectively).

**Figure 1.**
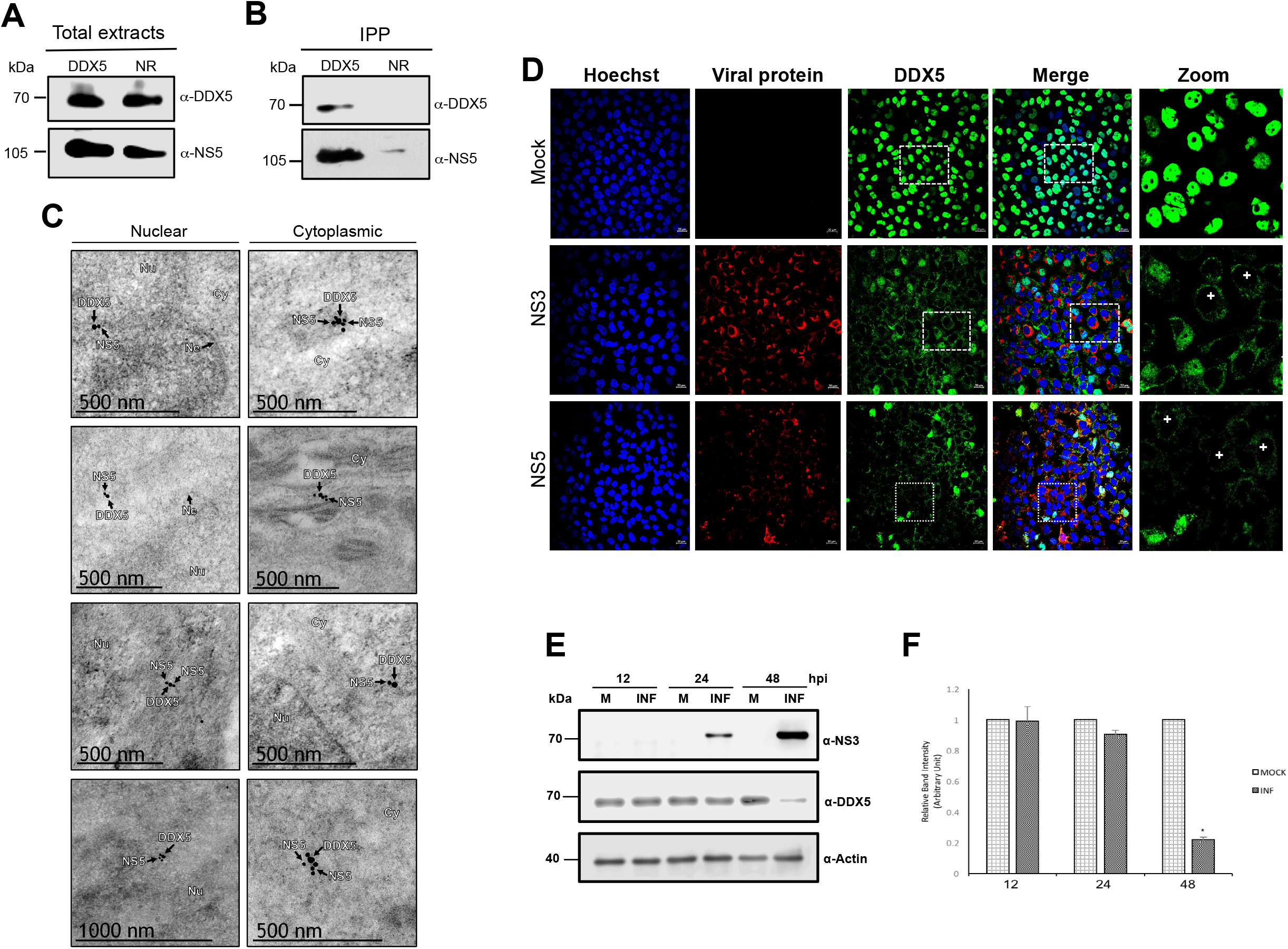
DDX5 protein interacts with NS5 and moves from nucleus to cytoplasm during DENV infection. (A) Total and (B) immunoprecipitated HuH-7 cell extracts with anti-DDX5 (DDX5) or anti-GFP (NR) antibodies were separated by SDS-PAGE and analyzed by western blot with anti-NS5 and anti-DDX5 antibodies. Representative western-blot assay from three independent experiments is presented. (C) Huh-7 cells mock-infected (Mock) or infected with DENV2 for 48 hrs, were incubated with an anti-DDX5 protein antibody and analyzed by confocal microscopy. Anti-NS3 and anti-NS5 proteins were used as controls of infection. Infected cells: (+). Nuclei were stained with Hoechst. (D) Huh-7 cells infected with DENV2 for 24 hrs were fixed and incubated with anti-NS5 (small gold particles) and anti-DDX5 (large gold particles) antibodies and analyzed by immunoelectron microscopy. (E) DDX5 protein levels were analyzed by western blot in total cell lysates obtained from mock (M) or DENV2 Huh7 infected cells at 12, 24, and 48 h. Representative images from three independent experiments are presented. (F) Graph representing DDX5 levels adjusted with β-actin and normalized with respect to mock-infected cells. Data are means ± standard error (S.E) of *n=3* independent experiments performed by duplicate. \**p<0.05.*

Since our main objective was to study new functions of NS5 protein in the nucleus, the NS5 protein, and the resin without His-NS5 (control) were incubated with nuclear cell extracts from mock-infected Huh-7 cells, and after four washes with different concentrations of NaCl and Imidazol, two elutions at 500 and 1000 mM of NaCl were obtained (Figure 2A and 2B, Supplementary material) Most of the proteins isolated interact specifically with His-NS5 (Figure 2A Supplementary material) since none of them were observed in the elution fraction of the negative control (Figure 2B Supplementary material). Mass spectrometry (Maldi-ToF) analysis of the eluted proteins from three independent experiments allowed the identification of 40, 42, and 36 proteins associated with His-NS5 that were absent in the proteins identified in the control resin without His-NS5. A total of 36 proteins that were identified in the 3 independent experiments were considered as true NS5-associated proteins (Table 1).

**Figure 2.**
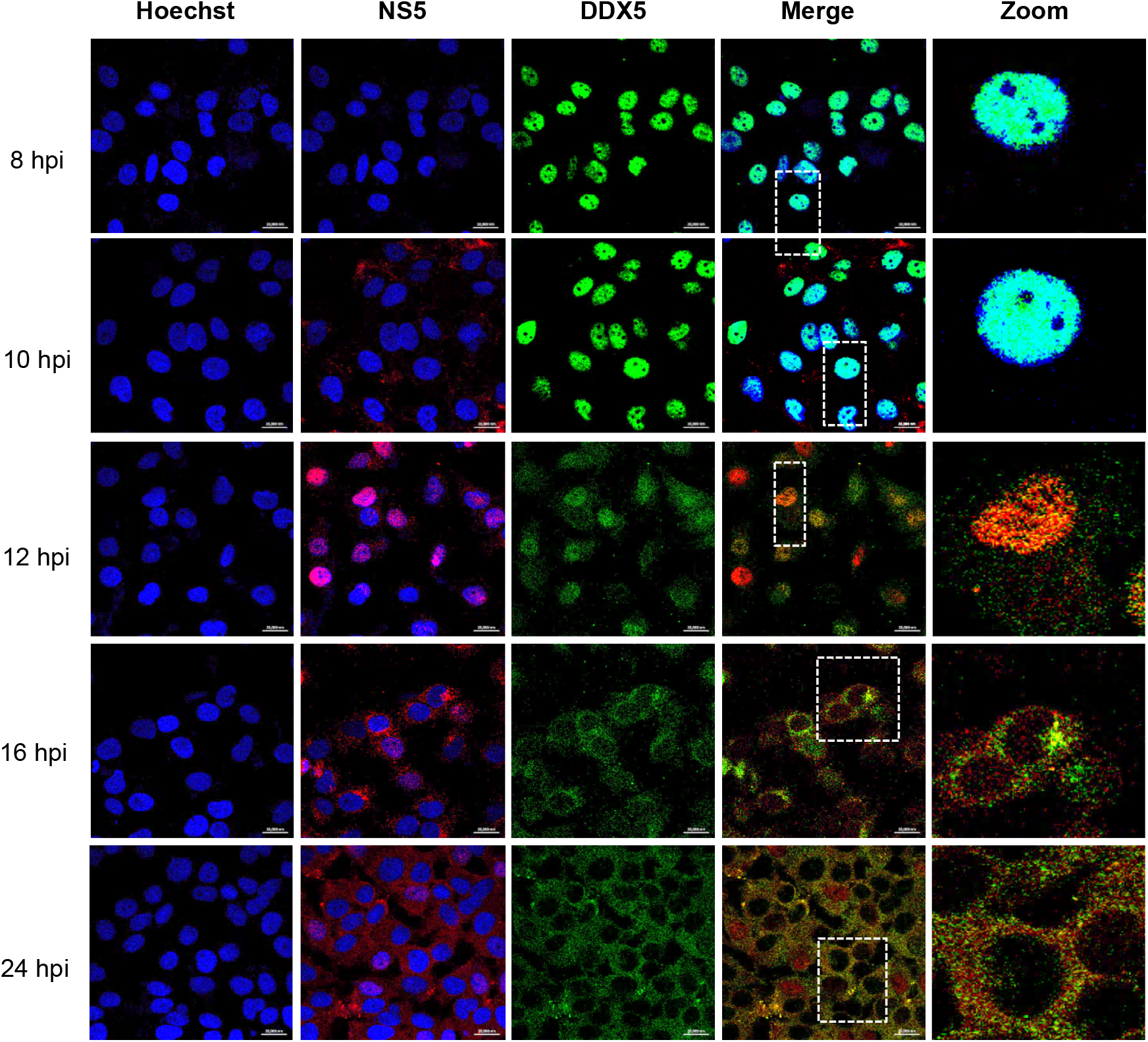
DDX5 is relocated from the nucleus to the cytoplasm after nuclear import of DENV2 NS5 protein. Huh-7 cells were infected with DENV2 for 8, 10, 12, 16, and 24 hrs and the subcellular localization of DDX5 and NS5 was analyzed by confocal microscopy using the monoclonal anti-DDX5 and the anti-NS5 protein antibodies respectively. Nuclei were stained with Hoechst. Representative images are presented.

### Functional characterization of NS5 interacting proteins

To further characterize the cell proteins identified, the functional categorization and subcellular localization were determined based on a Swiss-Prot TrEMBL database search http://www.uniprot.org/uniprot/P07437. The 36 cellular proteins identified are involved in seven biological processes: 6 out of the 36 (16.6 %) proteins are involved in translation; 5 (13.88%) in microtubule-based process; 5 (13.88%) are part of the spliceosome, 7 (19.44%) in metabolic process, 4 (11.11%) in stress response, 3 (8.33 %) in lipid metabolism, and 6 (15.6%) in other processes (Figure 3B Supplementary material). The 33% of the identified proteins were nuclear and 33% were cytoplasmic proteins; 17% were ribosomal proteins while only 11 and 6 % of the proteins were mitochondrial and cell membrane proteins respectively (Figure 3B, Supplementary material).

**Figure 3.**
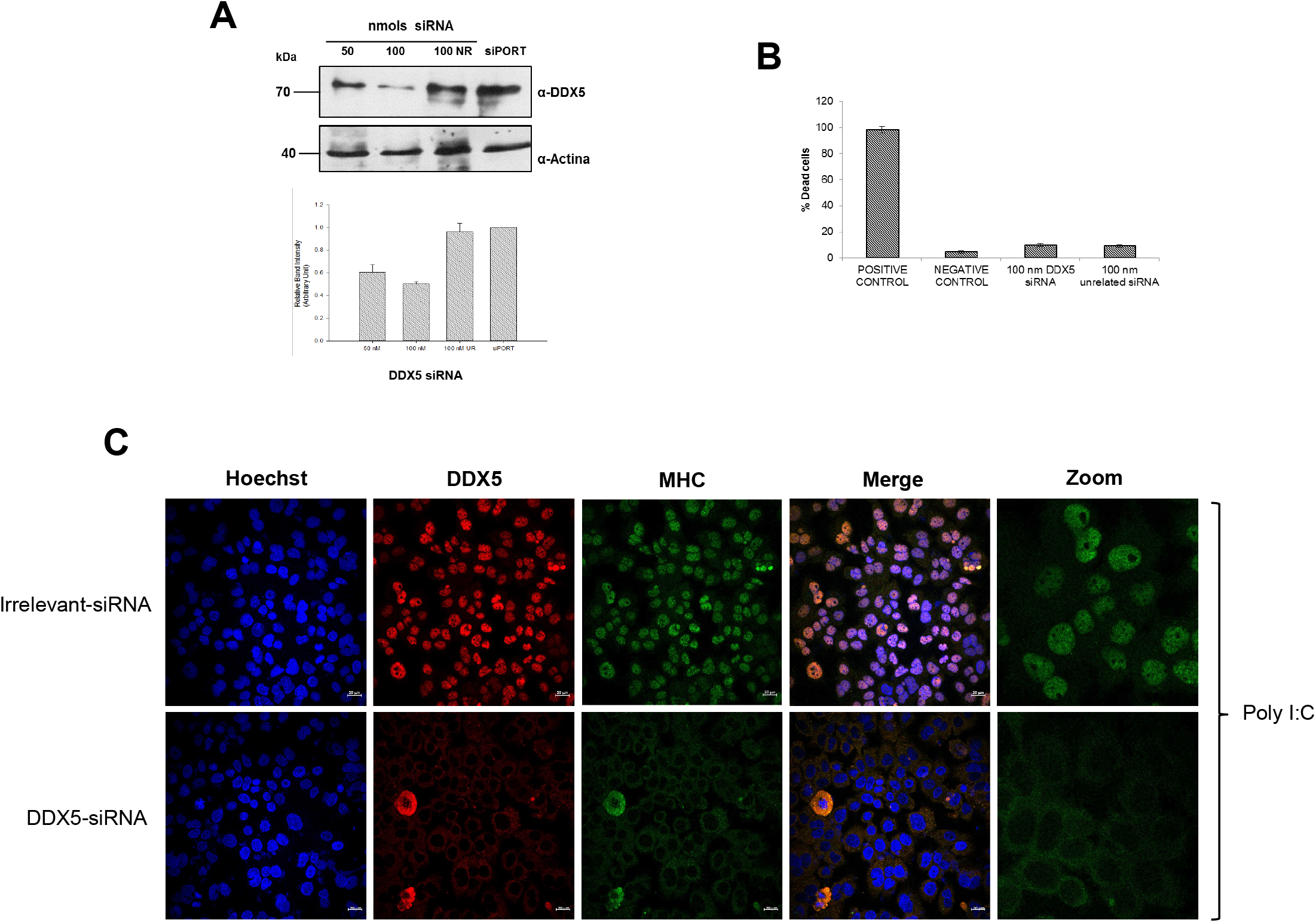
Silencing of DDX5 inhibits expression of MHC in response to poly I:C. Silencing of DDX5. (A) Cells were transfected with 50 and 100 nM of DDX5-siRNA or 100 nm of an irrelevant siRNA and the expression levels of DDX5 were determined by western blot. Anti-actin antibody was used as a loading control. Data are means ± standard error (S.E) of *n=3* independent experiments performed by duplicate. *\*p<0.05.* (B) Cell viability after silencing was determined by MTT assay. Data are means ± standard error (S.E) of *n=3* independent experiments performed by duplicate. *\*p<0.05.* (C) Huh-7 cells incubated with a DDX5-siRNA and with an irrelevant siRNA as a negative control, were treated with poly I:C and the expression of DDX5 and MHC was determined by confocal microscopy. Nuclei were stained with Hoechst. Representative images are presented.

Additionally, a protein network analysis of protein–protein interactions was performed to identify functionally linked proteins. The 36 cellular proteins identified were analyzed into STRING. Thirty-three out of the 36 proteins were found to be strongly linked either directly or indirectly through one or more interacting proteins suggesting the existence of functional linkages. Three out of the 36 identified proteins did not show any link at the chosen confidence level (STRING score=0.4). (Figure 3C, Supplementary material).

### DENV NS5 interacts with the DEAD-box RNA helicase DDX5 protein in Huh-7 cells

Considering that we were interested in identify nuclear proteins that interact with NS5 protein, we focused our attention in some proteins located in the nucleus such as the KH Type Splicing Regulatory Protein, the heterogeneous ribonucleoproteins K and F, insulin-like growth factor 2 mRNA binding protein 2, the splicing factor proline and the glutamine-rich, histon 2B, lamin A and the DEAD-box RNA helicase DDX5. Most of these proteins are involved in splicing or interact with RNA. Since it has been described that the DEAD-box RNA helicase DDX5 plays important roles in transcription modulation, differentiation and splicing^22^ and some other DDX proteins have been also identified as important factors during flavivirus infection^23–25^, we further study this protein and validate its interaction directly or indirectly with DENV NS5 protein by an immunoprecipitation assay. Thus, nuclear extracts from infected Huh-7 cells treated with micrococcal nuclease were incubated with the anti-DDX5 antibody and the immunoprecipitated fractions were analyzed by western blotting using specific anti-NS5 antibodies. The presence of the NS5 protein (105 kDa) was detected in the fractions immunoprecipitated with the anti-DDX5 (Figure 1B) but not when a non-related antibody (NR) was used (Figure 1B), suggesting the association between DDX5 and NS5, and validating the results obtained by the affinity chromatography assays. The presence of NS5 and DDX5 proteins in the nuclear extracts from total infected cells was confirmed by western blotting (Figure 1A). To further validate the association of NS5 with other proteins identified in the proteomic analysis, we selected the hnRNP F. As well as with the helicases, some other hnRNPs have been involved in DENV infection^26–28^. As well as it was observed with DDX5, NS5 was immunoprecipitated only in the presence of the hnRNP F antibodies and not in the presence of an unrelated antibody (Figure 4A, Supplementary material). The presence of NS5 (105 kDa) and hnRNP F proteins in the nuclear extracts from infected cells was also confirmed by western blotting (Figure 4B, Supplementary material).

**Figure 4.**
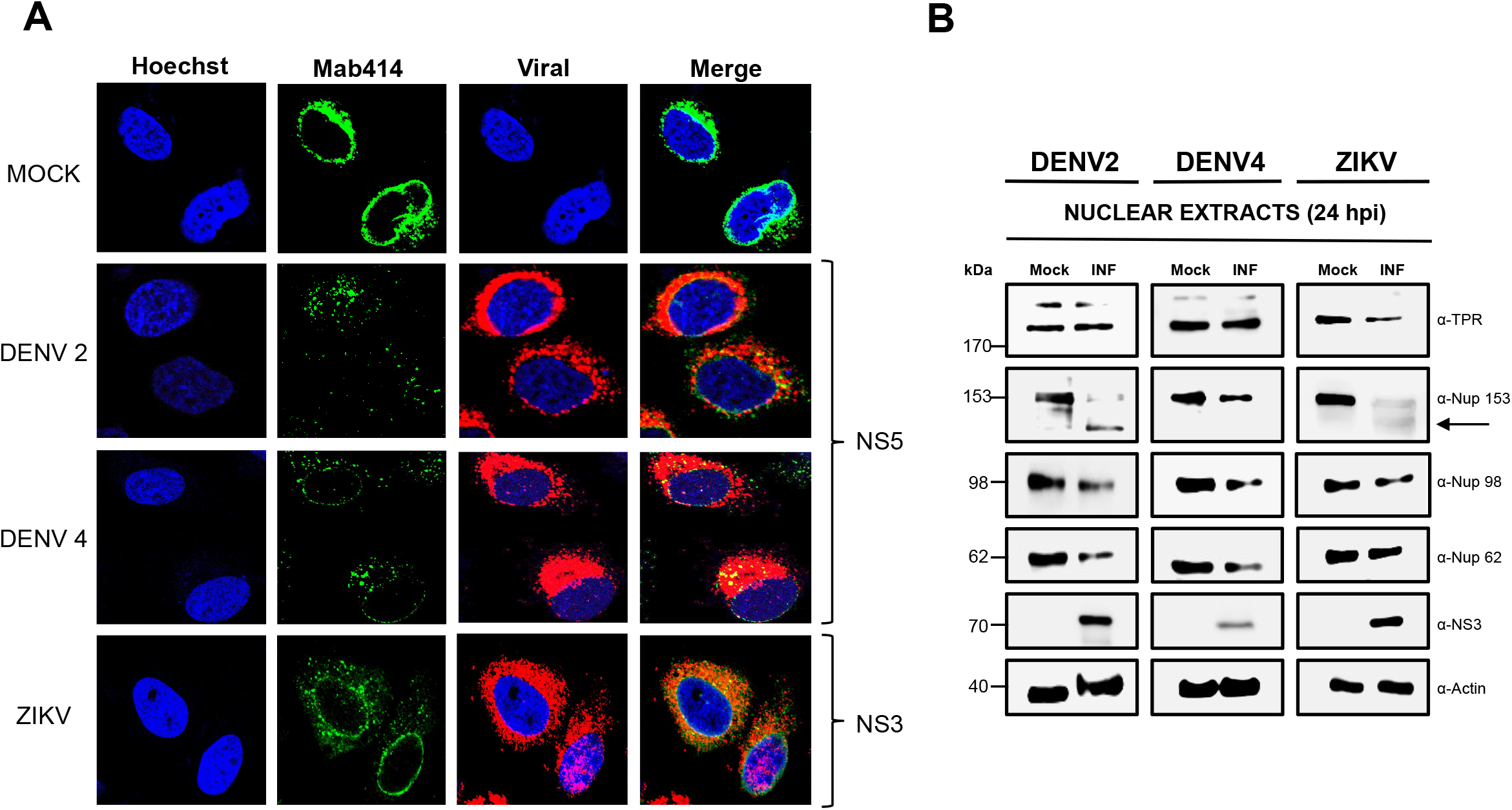
The integrity of the nuclear pore complex is altered after DENV and ZIKV infection. Huh-7 cells mock-infected (Mock) or infected with DENV2, DENV4 or ZIKV for 48 hrs, were incubated with the monoclonal antibody against the FG-rich sequence of nucleoporins, Mab414, and analyzed by confocal microscopy. Anti-NS3 and anti-NS5 proteins were used as controls of infection. Nuclei were stained with Hoechst. Representative images of three independent experiments are presented. (B) Levels of TPR, Nup153, Nup98, and Nup62 proteins were analyzed by western blot in whole cell lysates obtained from mock and DENV2, DENV4, and ZIKV infected cells for 24 hrs. Anti-NS3 antibody was used as a control of infection and anti-actin as a loading control. Representative western-blots assays of three independent experiments are presented.

### DEAD-box RNA helicase DDX5 protein changes its localization during DENV infection in Huh-7 cells

To confirm the nuclear location of DDX5, Huh-7 cells were infected with DENV2 at an MOI of 3 for 48 hrs, incubated with specific antibodies against DDX5 and analyzed by confocal microscopy, electron microscopy and western blotting. The first observation was that as expected, DDX5 was mainly located in the nucleus of mock-infected cells. Interestingly, during infection, DDX5 was relocated from the nucleus to the cytoplasm (Figure 1C). The association between DDX5 (big gold particles) and NS5 proteins (small gold particles) in the nucleus and the cytoplasm of infected cells was confirmed by electron microscopy (Figure 1D). Remarkably, the presence of DDX5 protein in the cytoplasm was strongly reduced at 48 hpi (Figure 1C) suggesting its degradation. To confirm this last result, the levels of DDX5 in total cell extracts from mock-infected (M) and DENV infected cells (INF) at 12, 24, and 48 hrs was analyzed by western blotting. As observed by confocal microscopy, the levels of DDX5 protein were reduced at 24 and 48 hrs post-infection (Figure 1E and 1F), confirming that DDX5 protein is degraded in the cytoplasm of DENV infected cells. A similar relocation from the nucleus to the cytoplasm was observed for the hnRNP F, another cell protein identified in our affinity chromatography assay that was confirmed to be associated with NS5 by immunoprecipitation assays. However, in this case, the protein is not degraded in the cytoplasm after infection (Figure 4A-E, Supplementary material).

To analyze in further detail the cytoplasmic relocation of DDX5, the subcellular localization of NS5 and DDX5 was analyzed at different times postinfection (Figure 2). As expected, at 8 and 10 hpi, DDX5 was mainly detected in the nucleus of infected cells while a discrete amount of NS5 is observed in the cytoplasm at 10 hpi. At 12 hpi, NS5 was clearly detected in the nucleus of infected cells, as has been described previously^29^; even though the majority of DDX5 is still in the nucleus, some of this protein is also observed in the cytoplasm. At 18 and 24 hpi, both DDX5 and NS5 were mainly in the cytoplasm of the infected cells, where they clearly colocalized (Figure 2). Considering all these results, three important questions arose, i) Why is DDX5 protein degraded? ii) How is a nuclear protein such as DDX5 now present in the cytoplasm? and iii) Which molecule is responsible for DDX5 degradation?

### DDX5 is involved in the immune response

The first obligated question about DDX5 is why this protein is degraded during DENV infection? One possibility is that DDX5 could be involved in the immune response against DENV infection. To analyze this possibility, DDX5 expression in Huh7 cells was silenced up to 50% using a specific DDX5-siRNA (Figure 3A and 3C), and then cells were treated with poly I:C and the expression of the major histocompatibility complex (MHC) was evaluated by confocal microscopy (Figure 3C).

As observed in cells treated with the irrelevant siRNA (NR), the incubation with poly I:C, caused an increased in the MHC expression, while in DDX5 silenced cells, the MHC expression was almost abolished (Figure 3C), supporting the idea that the degradation of DDX5 caused a significant inhibition in IFN response. Cell viability was not altered in the presence of the NR-and DDX5-siRNAs (Figure 3B).

This result correlates well with the suggested function of DDX5 and other DDX helicases in the innate immune response during viral infections^23, 30^

### The infection with DENV and ZIKV induces cleavage/degradation of some nucleoporins of the nuclear pore complex

Our next important question was how a nuclear protein such as DDX5 reaches the cytoplasm of infected cells? Besides DDX5 and hnRNP F the relocation of other nuclear proteins such as PTB, La, hnRNP K, DDX21, and HMGB1 to the cytoplasm during DENV infection has also been described^23, 26, 31–34^ Moreover, an important reduction in the transport of mature mRNAs from the nucleus to the cytoplasm has also been described in DENV infected cells^14^. All these results suggest that flavivirus infection induces modifications in the nuclear-cytoplasmic transport. The nuclear-cytoplasmic transport is regulated by the nuclear pore complex (NPC) and by the transport factors, like karyopherins called importins, which transport proteins and RNAs from the cytoplasm to the nucleus, and exportins to exit the nucleus. The NPC is composed of several copies of approximately 30 different proteins, called nucleoporins (NUPs). Most of the NUPs are structural proteins; however, the NUPs that form the central channel of the NPC harbor a Phe and Gly-rich repeats (FG-NUPs) that play an important role in the transport of cargo molecules through the NPC. For this reason, and considering that in DENV infected cells, several nuclear proteins are translocated to the cytoplasm and the mature mRNAs transport to the cytoplasm is inhibited^14^, we initially analyzed the integrity and function of some of the components of the NPC. The distribution and integrity of the FG-NPC were evaluated using the monoclonal antibody Mab414, which is directed to the FG repeats present in several NUPs (Figure 4A). Staining of a ring around the nucleus was clearly observed in mock-infected cells incubated with the Mab414; however, this ring was disassembled in cells infected with the two DENV2 and DENV4 serotypes, supporting the idea that DENV infection alters the integrity or distribution of the NUPs that contain FG repeats (Figure 4A). Given the close relationship between DENV and ZIKV, staining of the NPC with the same antibody was performed in cells infected with ZIKV. A partial disruption of the ring structure and an important reduction in the intensity of this structure was detected in cells infected with ZIKV, suggesting that the integrity or distribution of the NUPs harboring FG repeats is modified during infection with both flaviviruses (Figure 4A).

Since Mab414 detects several NUPs harboring the FG repeats, the next step was to analyze the integrity of specific NUPs that are involved in RNA and protein transport through the NPC by western blotting. Interestingly, at 24 hpi with DENV2 and DENV4, an important reduction in the amount of Nup153, Nup98, and Nup62 but not of TPR was observed (Figure 4B). In contrast, at 24 hpi with ZIKV, a significant reduction in the amount of TPR, Nup153, and Nup98 but not of Nup62 was observed (Figure 4B). These results strongly suggest that DENV and ZIKV infection induced the cleavage/degradation of some NUPs involved in nuclear-cytoplasmic transport, being only Nup153 and Nup98 altered in both DENV and ZIKV infection.

Since it has been described that Nup98 and Nup62 play important roles in nuclear-cytoplasmic transport of mRNAs and some proteins, we analyzed in further detail the integrity of these two NUPs during the infection of DENV and ZIKV. Using an anti-Nup62 specific antibody, we detected a homogenous distribution of this protein in the nucleoplasm of mock-infected cells (Figure 5A). However, in DENV2 and DENV4 infected cells a significant reduction of the protein in the nucleoplasm was observed at 48 hpi, with no presence in the cytoplasm (Figure 5A). On the other hand, a relocalization of Nup62 from the nucleus to the cytoplasm of the ZIKV infected cells was clearly observed, in comparison to the nuclear localization of Nup62 in mock-infected cells (Figure 5A). Moreover, the abundance of this nucleoporin was not altered. To confirm these results, the expression levels of Nup62 during DENV and ZIKV infection were analyzed by western blotting at different times post-infection. A slight reduction of 22 and 31% in the expression levels of Nup62 was detected at 24 hpi in cells infected with DENV2 and DENV4 respectively. Moreover, the reduction in the expression levels of Nup62 at 48 hpi was more prominent (52 and 71% respectively) (Figure 5B). In concordance with the images observed by confocal microscopy, the abundance of Nup62 in cells infected with ZIKV was not altered at 12, 24, and 48 hpi (Figure 5B).

**Figure 5.**
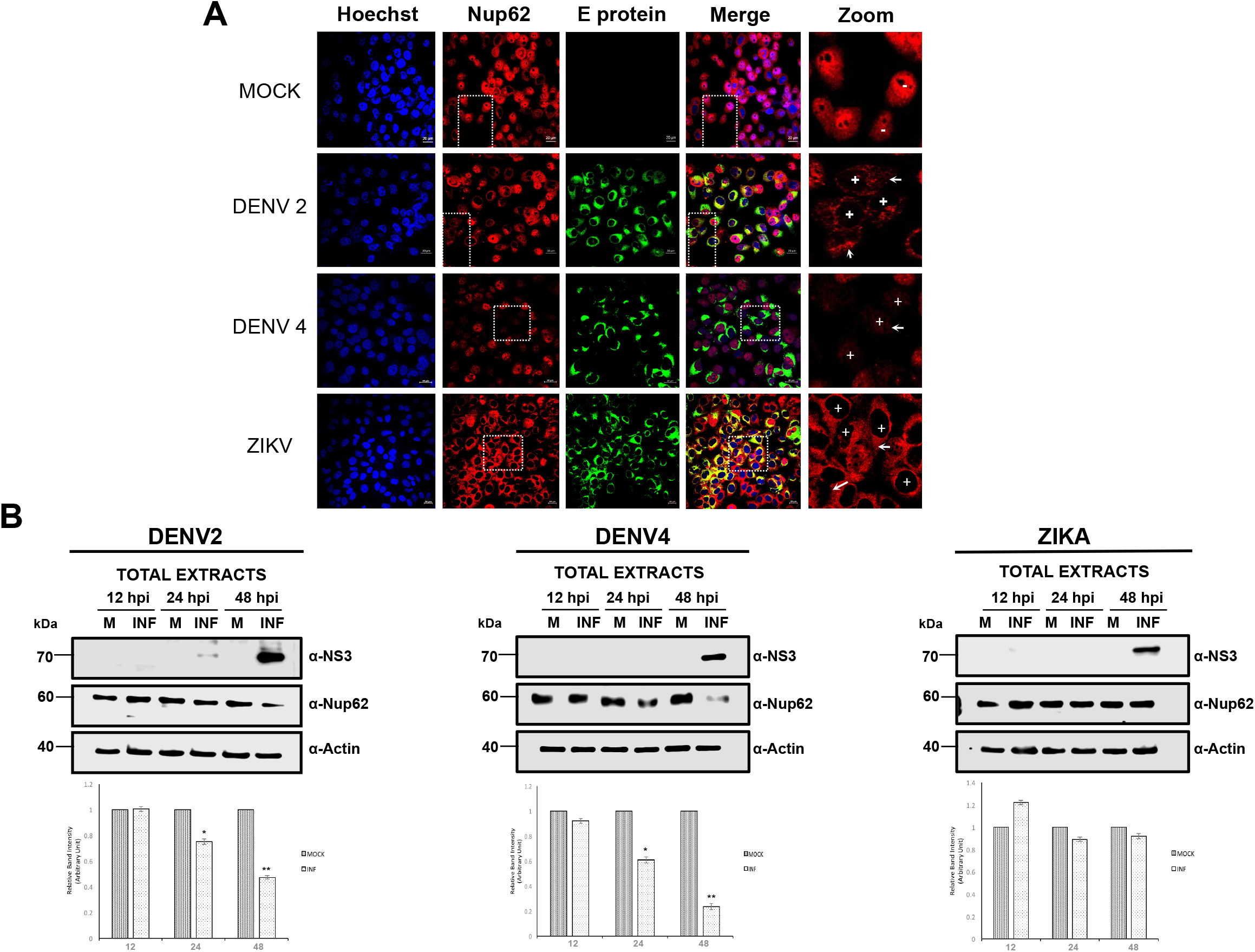
The integrity and location of the Nup62 is altered during DENV and ZIKV infection. Huh-7 cells mock-infected (Mock) or infected with DENV2, DENV4, or ZIKV for 48 hrs, were incubated with a monoclonal anti-Nup62 antibody and analyzed by confocal microscopy. The anti-E protein antibody was used as control of infection. Nuclei were stained with Hoechst. Representative images of three independent experiments are presented. (B) Levels of Nup62 protein were analyzed by western blot in whole cell lysates from mock or DENV2, DENV4, and ZIKV infected cells for 12, 24, and 48 hrs. Anti-NS3 antibody was used as a control of infection and anti-actin as a loading control. Representative western-blots assays from three independent experiments are presented. Graph represents Nup62 protein levels compared with β-actin. Data are means ± standard error (S.E) of *n=3* independent experiments performed by duplicate. *\*p<0.05.*

When the integrity and abundance of Nup98 during DENV2, DENV4, and ZIKV were analyzed by confocal microscopy at 24 hrs post-infection using a specific anti-Nup98 antibody, the ring structure around the nucleus observed in mock-infected cells was clearly disrupted in cells infected with all these three viruses, supporting the idea that an important disruption in the distribution of Nup98 occurs in infected cells (Figure 6A). While in DENV2 and DENV4, the anti-Nup98 antibody signal was observed partially in the cytoplasm, no signal was observed in the ZIKV infected cells (Figure 6A). To further analyze the effect of DENV and ZIKV infection in the integrity of the Nup98, a western blotting assay was performed at 12, 24, and 48 hpi. A significant reduction in Nup98 protein levels was detected at 24 hpi with DENV2, DENV4, and ZIKV (71, 52 and 83% respectively) (Figure 6B). This reduction was more dramatic at 48 hpi mainly for ZIKV (82, 40 and 91% for DENV2, DENV4 and ZIKV respectively), indicating that Nup98 is cleaved/degraded at 48 hpi with DENV and ZIKV. In summary, while DENV induces a delocalization and a significant reduction in the expression levels of Nup62 and Nup98, ZIKV induced a delocalization of Nup62 and a much more evident reduction in the levels of Nup98.

**Figure 6.**
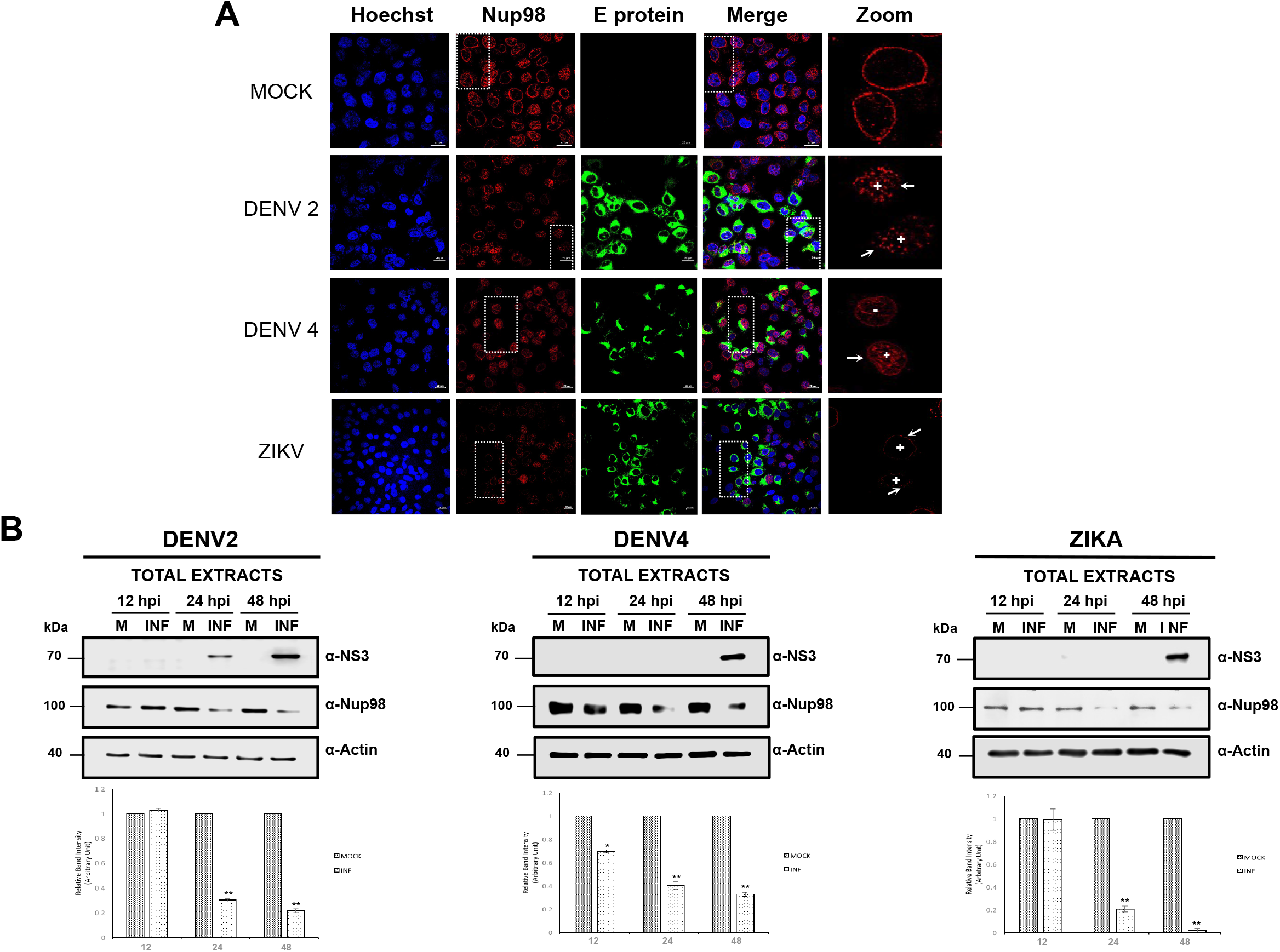
The integrity and location of the Nup98 is altered during DENV and ZIKV infection. Huh-7 cells mock-infected (Mock) or infected with DENV2, DENV4, or ZIKV for 48 hrs, were incubated with a monoclonal anti-Nup98 antibody and analyzed by confocal microscopy. The anti-E protein antibody was used as control of infection. Nuclei were stained with Hoechst. Representative images of three independent experiments are presented. (B) Levels of Nup-98 protein were analyzed by western blot in whole cell lysates from mock or DENV2, DENV4, and ZIKV infected cells for 12, 24, and 48 hrs. Anti-NS3 antibody was used as a control of infection and anti-actin as a loading control. Representative western-blots assays of three independent experiments are presented. Graph represents Nup-62 levels comparing with β-actin. Data are means ± standard error (S.E) of *n = 3* independent experiments performed by duplicate. *\*p<0.05.*

### A serine protease is responsible for NUPs cleavage/degradation in DENV and ZIKV infected cells

Thus, the next question to answer was which molecule could be involved in Nup98 and Nup62 cleavage/degradation during flavivirus infection. Since DENV and ZIKV encode for a serine-protease (NS2B-NS3), the first approach was to analyze the effect of two serine-proteases inhibitors, Leupeptin and TLCK.HCl in Nup62 and Nup98 cleavage during DENV infection (Figure 5A Supplementary material). Both inhibitors were able to prevent NUPs cleavage (Figure 5A Supplementary material), suggesting that a serine-protease such as NS2B-NS3 could be involved in Nup62 and Nup98 degradation. To support this possibility, a prediction of the possible fragments generated by NS3 activity of both viruses in Nup62 and Nup98 proteins was obtained using the software Prosite (Figure 5B Supplementary material). The two predicted cleavage products (cp) from Nup62 coincide with the ones observed in the SDS-PAGE obtained from cells incubated in the absence of both inhibitors (VC) (marked as cp1 and cp2) as well as with one of the predicted cleavage products from Nup98 (cp1) (Figure 5A Supplementary material). The solvent-exposed RRK sequence (Arginine-Arginine-Lysine) of Nup98 in the position 777-780, support the idea that NS2B-NS3 could be involved in NUPs degradation. All these results taken together suggest that NS3 or a cellular serine-protease induced during DENV infection could be involved in the NUPs degradation.

### NS3 localizes in the perinuclear region and in the nucleus of the infected cells at early times post-infection and it is responsible for FG-NUPs degradation

To analyze the possible participation of NS3 in NUPs cleavage/degradation, cells were transfected with a plasmid encoding the NS2B-NS3 of DENV2 and ZIKV and the distribution and integrity of the FG-NUPs were analyzed by confocal microscopy using Mab414 antibody (Figure 7). While in mock-transfected cells the Mab414 antibody stained a ring structure around the nucleus, at 24 hpt a partial disruption of the nuclear ring structure was observed, that was much more pronounced at 48 hpi suggesting that the NS2B-NS3 is responsible for the FG-NUPs degradation (Figure 7).

**Figure 7.**
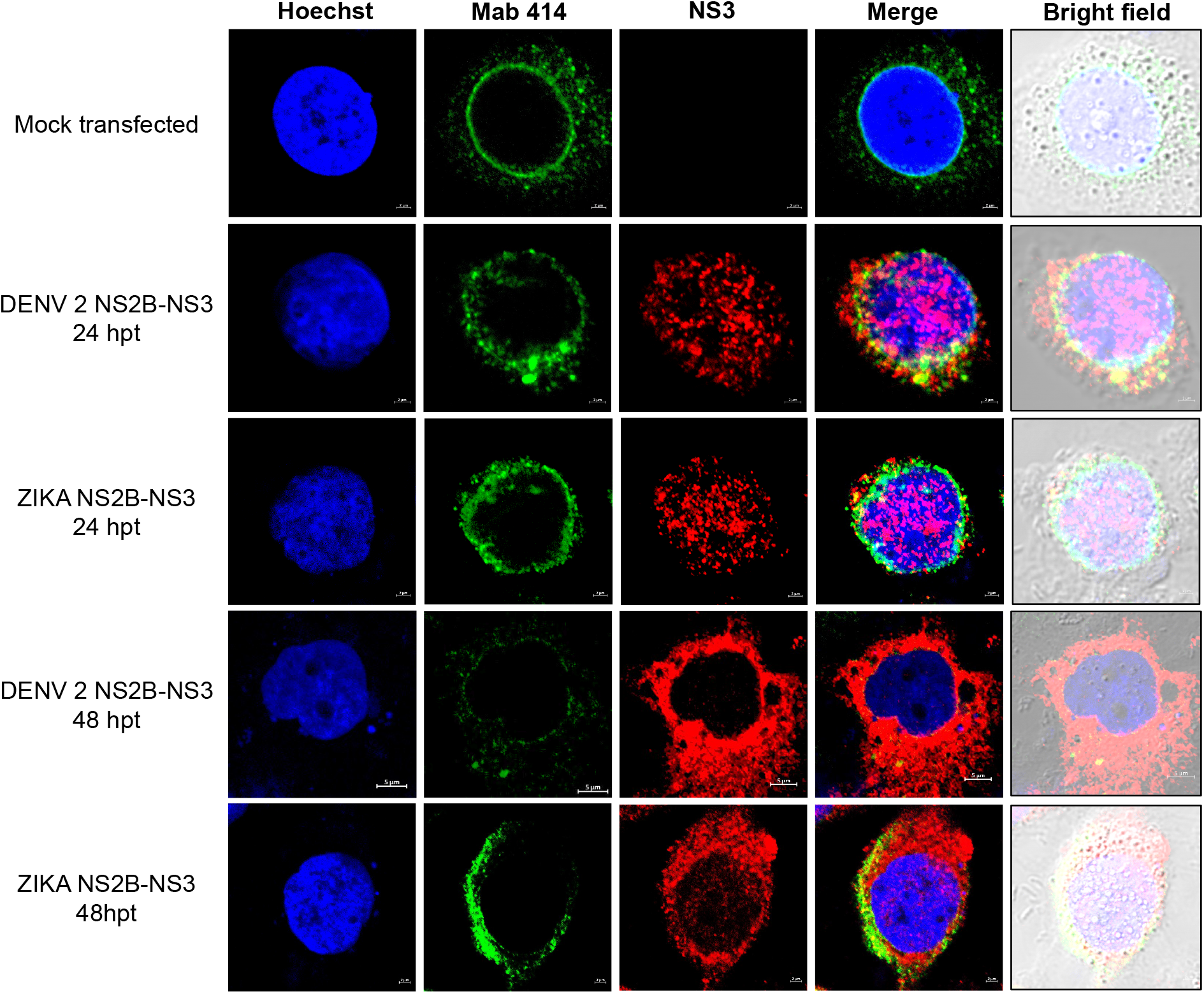
Transfection of NS2B-NS3 proteins from DENV2 and ZIKV induce disruption of FG-rich sequence nucleoporins. Huh-7 cells mock tranfected or transfected for 24 and 48 hrs with DENV2 or ZIKV NS2B-NS3 proteins were incubated with the anti-FG-rich sequence nucleoporins antibody (Mab-414) and with the anti-NS3 protein antibody and the integrity and subcellular localization of NS3 protein were analyzed by confocal microscopy. Nuclei were stained with Hoechst. Representative images of three independent experiments are presented

To further confirm the participation of the protease activity of NS2B-NS3 in the FG-NUPs degradation, cells were transfected with the NS2B-NS3 mutant S135A, which lacks the protease activity (Figure 8). The expression of this mutant protease did not alter the distribution and integrity of the FG-NUPs, confirming that the protease activity of NS2B-NS3 is responsible of the FG-NUPs degradation (Figure 8).

**Figure 8.**
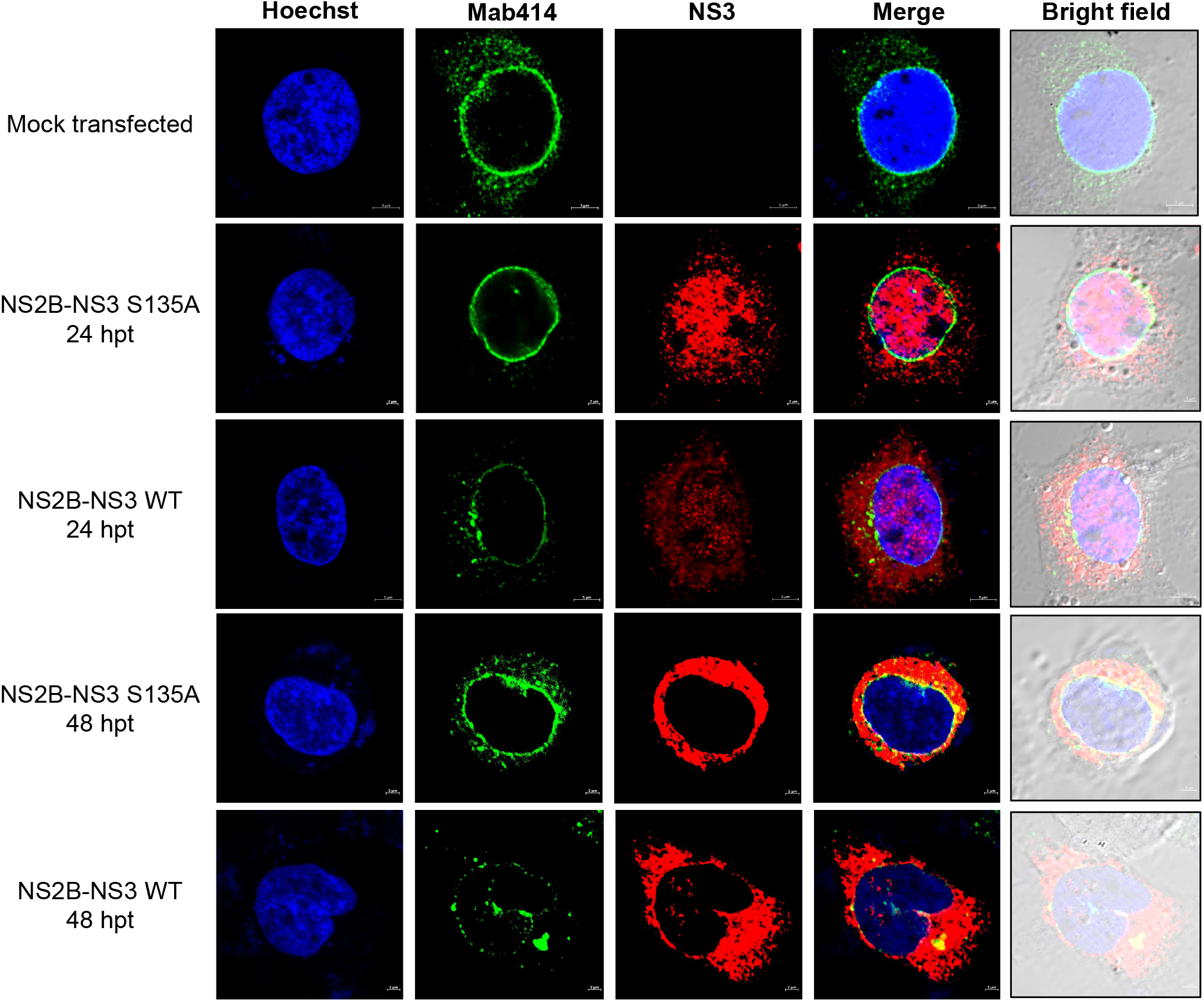
Transfection of the wild type but not with the mutant NS2B-NS3 protein from DENV2 induces disruption of FG-rich sequence nucleoporins. Huh-7 cells transfected for 24 and 48 hrs with wild type and DENV2 mutant NS2B-NS3 protein. Nucleoporins integrity and subcellular localization of NS3 were analyzed by confocal microscopy using the Mab-414 and the anti-NS3 protein antibody respectively. Nuclei were stained with Hoechst. Representative images of three independent experiments are presented.

It has been widely reported that degradation of nucleoporins occurs by the activity of viral proteases located in the perinuclear and the nuclear regions, such as the 2A and 3CD from rhinovirus^35, 36^ Nup98 is degraded early after DENV and ZIKV infection and it is located in both the nuclear and the cytoplasmic regions of the NPC; thus, it is possible that its degradation may be mediated by the protease present in the perinuclear region or in the nucleus of the infected cells. To analyze this possibility, Huh-7 cells were infected with DENV2, DENV4, and ZIKV and the subcellular location of NS3 at different times post-infection was analyzed by confocal microscopy (Figure 9). Interestingly, in cells infected with DENV2 and DENV4, the NS3 protein was detected in the nucleus and in the cytoplasm of the infected cells at 8 hpi, while in cells infected with ZIKV the same distribution of NS3 was observed at 12 hpi. NS3 protein from DENV2 and ZIKV was observed mainly in the perinuclear region and in the cytoplasm from 16 to 24 hpi, which is the location that has been reported up to now for NS3 in infected cells (Figure 9). These results are in accordance with the subcellular localization of both the mutant and the wt NS2B-NS3 transfected proteins in the nucleus and the cytoplasm at 24 hpt and in the cytoplasm at 48 hpt. These results indicate that NS3 protein from DENV and ZIKV is present in close proximity to the NPC early after transfection/infection and suggest that it may be a shuttling protein in both transfected and infected cells (Figures 7–9).

**Figure 9.**
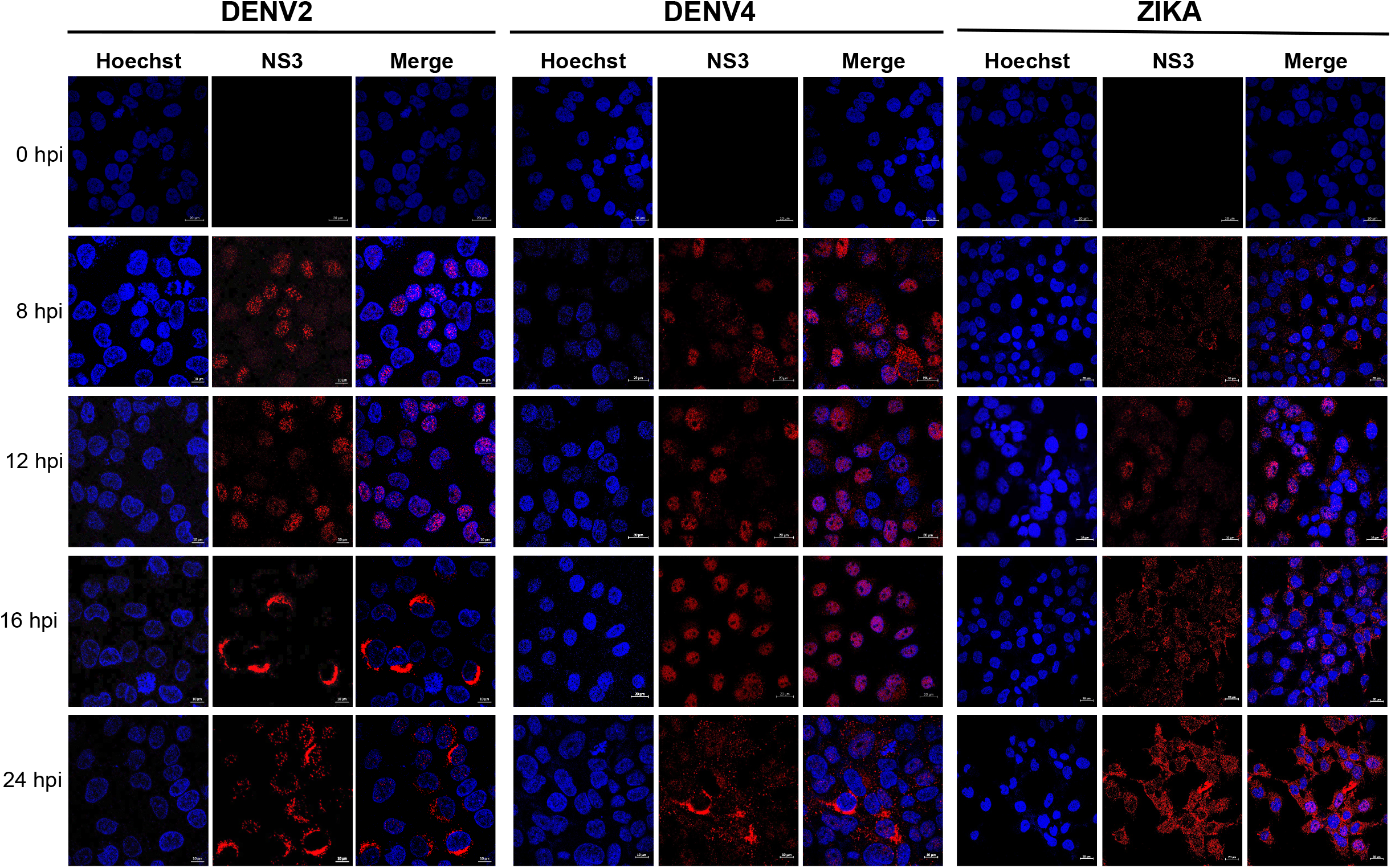
NS3 protein is located in the nucleus and perinuclear region early after DENV, and ZIKV infection. Huh-7 cells infected for 0, 8, 12, 16, and 24 hrs with DENV2, DENV4, or ZIKV were incubated with anti-NS3 antibody and analyzed by confocal microscopy. Nuclei were stained with Hoechst. Representative images of three independent experiments are presented.

### The degradation of some FG-NUPs alters mRNA export

So far, our results indicated that during DENV and ZIKV infection, a disruption of FG-NUPs by the action of NS2B-NS3 occurs, suggesting that the nuclear-cytoplasmic transport might be altered. Specifically, it has been described that Nup98, which is cleaved by both viruses very early after infection, is one of the NUPs that together with Nup153 and TPR, also altered in DENV and ZIKV infection (Figure 2), are involved in mature mRNA export^37^. Thus, to determine if the nuclear-cytoplasmic transport is altered, the location of mRNA in Huh-7 cells infected with DENV or transfected with DENV NS2B-NS3 was analyzed using oligo dT coupled to Cy5 probe in an *in situ* hybridization assay (Figure 10). As expected, in mock-infected cells, most of the poly A tailed mRNA is present in the cytoplasm, while a dramatic reduction of poly A tailed mRNA in the cytoplasm is observed in infected cells. A similar reduction in the amount of poly A tailed mRNA was also observed in the cells transfected with the NS2B-NS3 (Figure 10), suggesting that in both cases, the mRNAs are not exported to the cytoplasm. Interestingly, when the mutant NS2B-NS3 protease was transfected, the amount of poly A tailed mRNAs in the cytoplasm was similar than the observed in the mock-infected cells. All these results support the idea that the degradation of some FG-NUPs such as Nup98 induced by NS2B-NS3 alters dramatically the mRNA export through the NPC.

**Figure 10.**
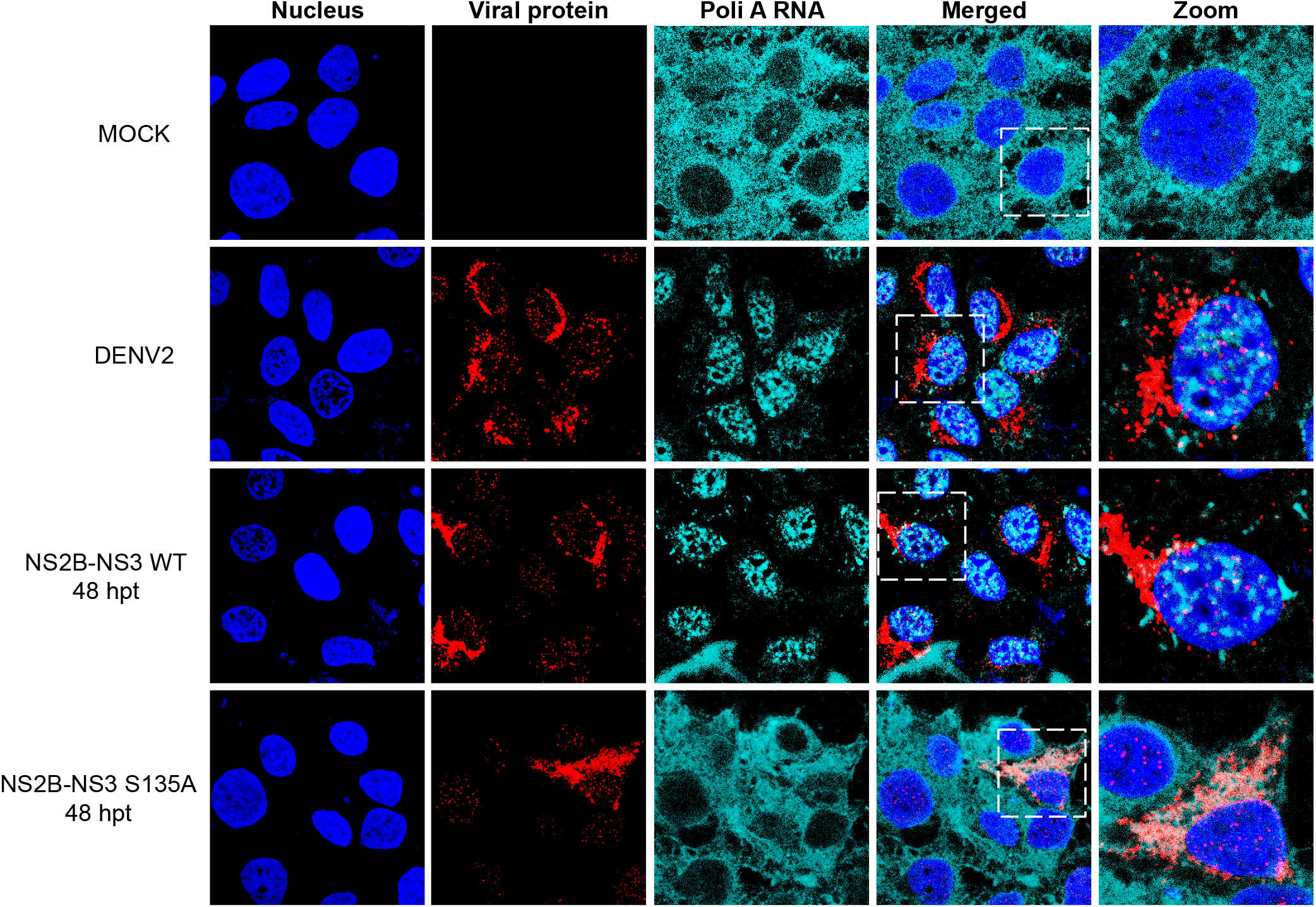
Transfection of the wild type but not the mutant NS2B-NS3 protein from DENV2 inhibits poly-A tailed mRNA export. Huh-7 cells were transfected for 48 hrs with the DENV2 wt- and mutant NS2B-NS3 proteins and the subcellular localization of poly A RNAs and NS3 was determined using oligo-dT coupled to Cy5 and anti-NS3 protein antibody respectively, using a confocal microscope. Nuclei were stained with Hoechst. Representative images of three independent experiments are presented.

### NS5 is required for the relocation of nuclear proteins to the cytoplasm

Considering that one of the main consequences of the NUPs degradation was the dramatic reduction in mRNA export, we wonder if NUPs disruption caused by NS2B-NS3 could also be related to the cytoplasmic localization of nuclear proteins such as DDX5 and hnRNP F. Surprisingly, as observed by confocal microscopy (Figure 11), after transfection with the DENV active protease, neither DDX5 nor hnRNP F were relocated to the cytoplasm (Figure 11) as during infection (Figure 1), indicating that other components present during infection are required for the presence of these proteins in the cytoplasm.

**Figure 11.**
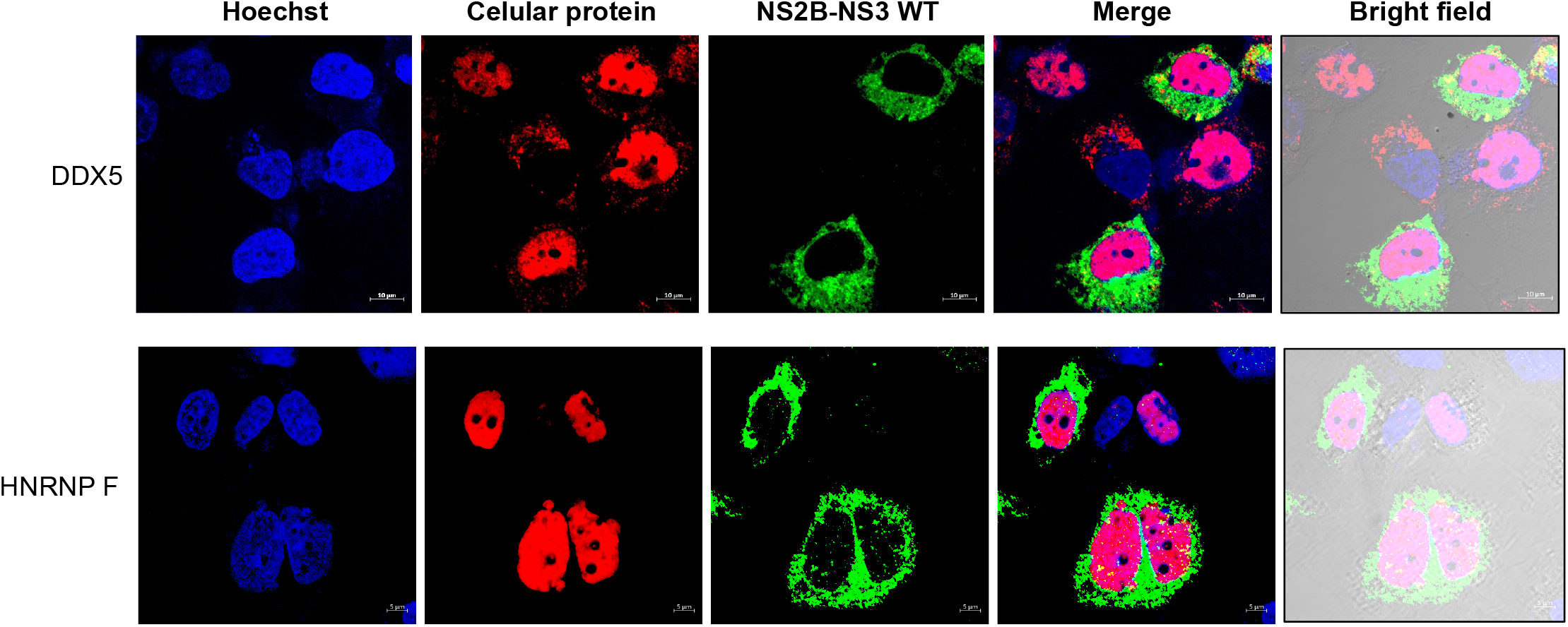
Export of DDX5 and hnRNP F proteins is not altered by DENV2 NS2B-NS3 protein transfection. Huh-7 cells transfected for 48 hrs with wt DENV2 NS2B-NS3 protein were incubated with anti-NS3, DDX5, or hnRNP F antibodies and subcellular localization of all proteins were analyzed by confocal microscopy. Nuclei were stained with Hoechst. Representative images of three independent experiments are presented.

Given that DDX5 and hnRNP F were not relocated to the cytoplasm after transfection of NS2B-NS3, we wonder if NS5, which interacts with both proteins, could be responsible to their cytoplasmic location. To analyze this possibility, Huh7 cells were transfected with a plasmid, which encodes the DENV2 NS5 protein fused to HA tag and the localization of DDX5 and hnRNP F was evaluated at 24 and 48 hpt by confocal microscopy (Figure 12). At 24 and 48 hpt both DDX5 and hnRNP F proteins are present in the cytoplasm (Figures 12A, and 12B respectively) in contrast with their nuclear location in the mock-transfected cells, confirming that NS5 is required for the relocation of nuclear proteins such as DDX5 and hnRNP F to the cytoplasm.

**Figure 12.**
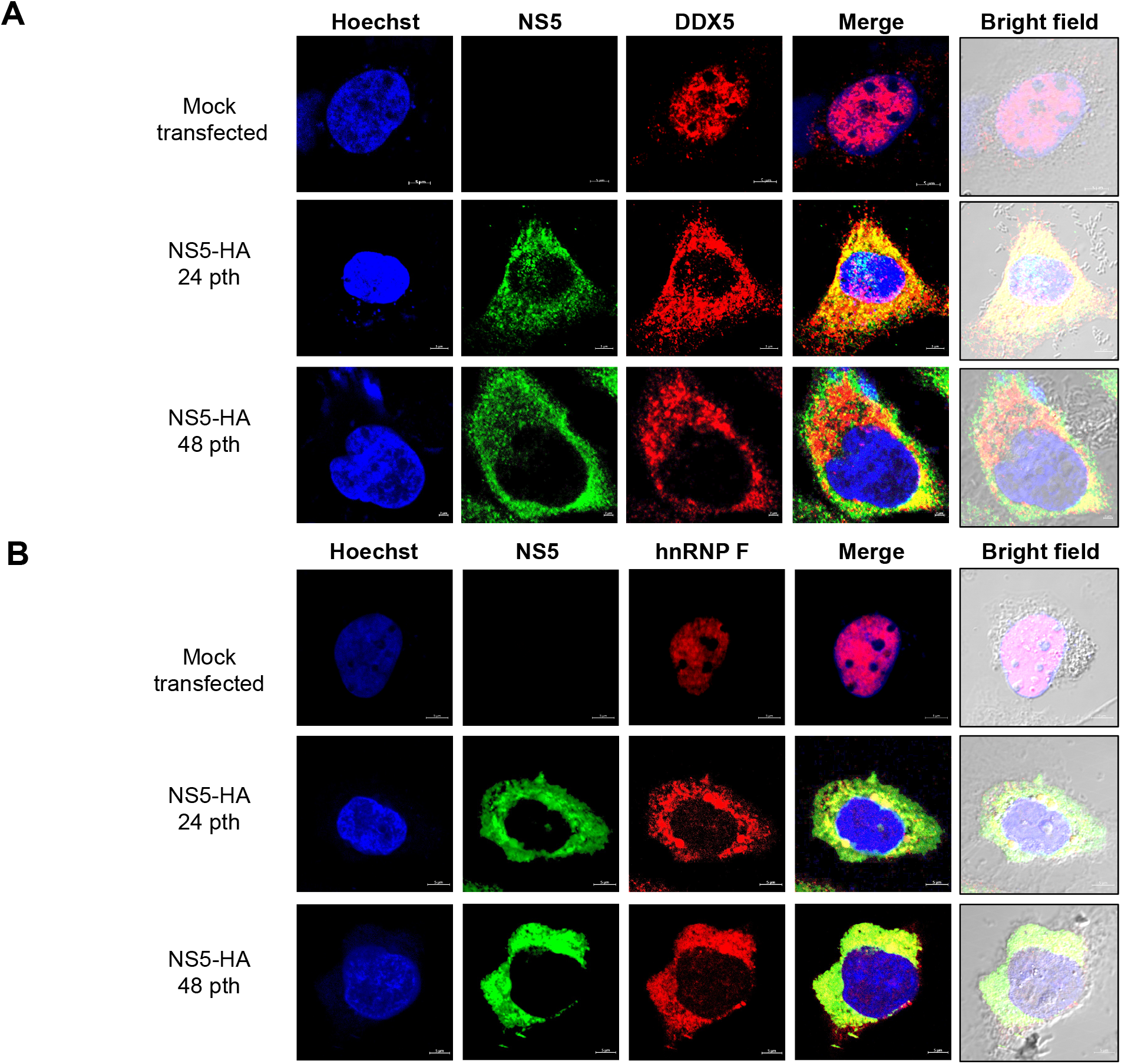
Transfection of DENV2 NS5 induces export of DDX5 and hnRNP F proteins. Huh-7 cells were transfected with NS5 protein from DENV2 for 24 and 48 hrs and the subcelular localization of DDX5, hnRNP F, and NS5 was analyzed by confocal microscopy using the anti-DDX5, anti-hnRNP F, and anti-NS5 protein antibodies respectively. Nuclei were stained with Hoechst. Representative images are presented.

While transfection of NS5 confirmed its role in the relocation of DDX5 and hnRNP F, the molecule involved in DDX5 degradation shown at 48 hpi remained undetermined. To analyze the possible role of the serine-protease NS2B-NS3 in DDX5 degradation, Huh-7 cells were transfected with NS5 and with NS2B-NS3 proteins and the integrity and location of DDX5 was analyzed by confocal microscopy. As observed in Figure 13, the transfection of both NS5 with NS2B-NS3 but not with NS2B-NS3 S135A was sufficient to induce translocation and degradation of DDX5 as in infected cells (Figure 13A), indicating that the DDX5 translocated to cytoplasm by NS5 is degraded by NS2B-NS3 after infection. These results correlate with the two putative serine-protease cleavage sites in DDX5 as revealed by the *in silico* analysis (Figure 13B).

**Figure 13.**
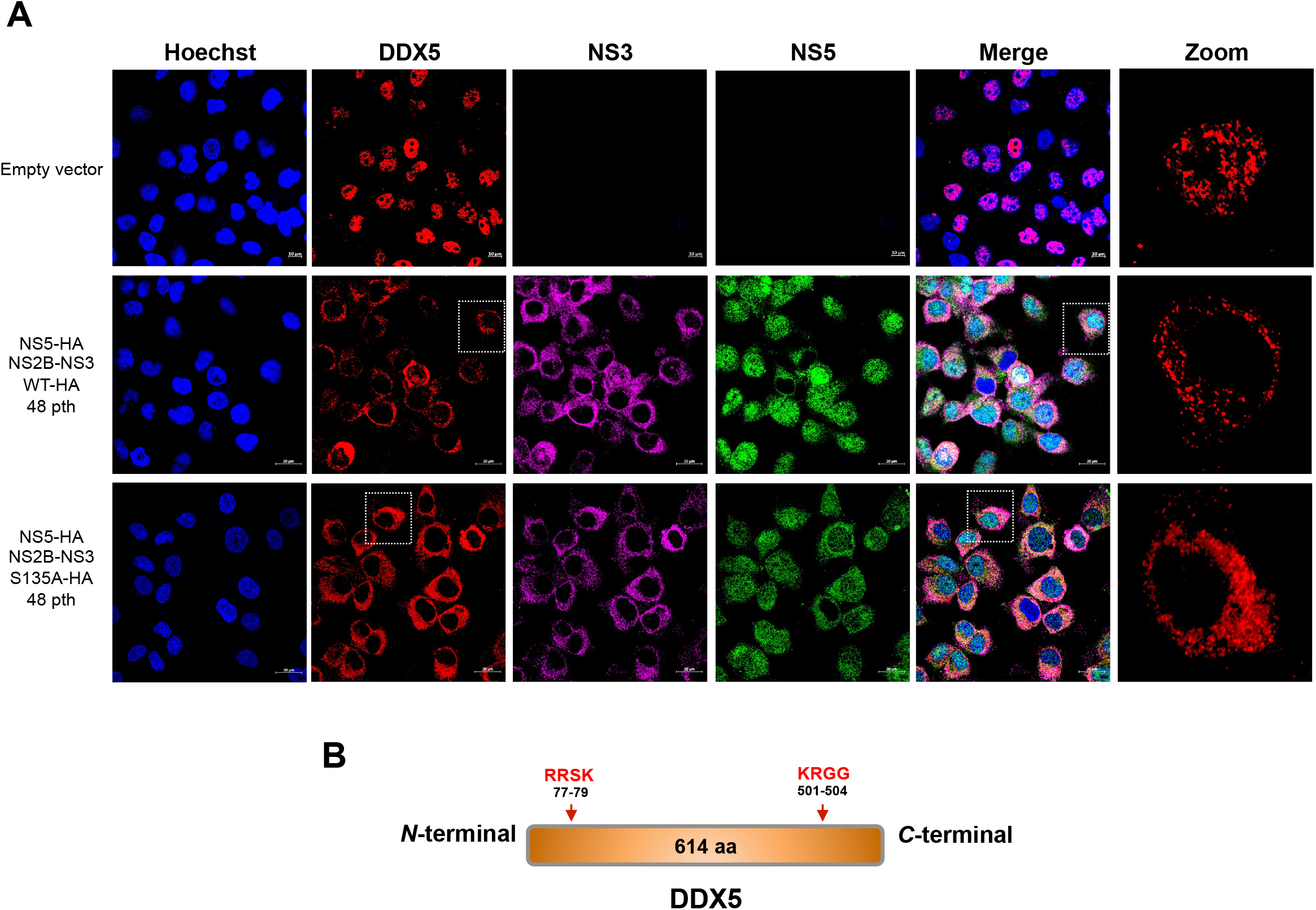
NS5 and NS2B-NS3 proteins from DENV2 induces export and degradation of DDX5. (A) Huh-7 cells were transfected with NS5 protein, and with the wild type or mutant NS2B-NS3 from DENV2 for 48 hrs. The subcellular localization of DDX5, NS5, and NS3 proteins was analyzed by confocal microscopy. Nuclei were stained with Hoechst. Representative images are presented. (B) Predicted targets of NS2B-NS3 serin-protease in the protein sequence of DDX5.

All these results presented here support the idea that the two viral proteins with enzymatic activity, NS3, and NS5, are involved in the induction of important changes in the nuclear-cytoplasmic transport during DENV and ZIKV infection. These changes favored the control of immune response and a significant reduction in the amount of mRNA to be translated in the cytoplasm favoring an advantageous cellular environment crucial for viral replication and immune response modulation.

## Discussion

Given the limited number of genes encode for many viruses, each viral protein has to perform several functions to manipulate cellular proteins, components and processes to favor its own replication in the host cell. It is well known that during flavivirus infection cytoplasmic cell proteins, components and organelles are hijacked; however, the manipulation of nuclear components and its consequences are unknown. In this respect, although flaviviral replicative cycle takes place in the cytoplasm, two viral proteins, C and NS5, are also present in the nucleus of the infected cells.

Due to the important functions of NS5 protein in the flavivirus replicative cycle, and the striking presence of this protein in the nucleus, different groups have used several strategies to isolate and identify cellular proteins that interact with NS5. By affinity chromatography assays with a recombinant NS5 protein, our group identified 36 proteins associated with NS5. The association of NS5 with five of them: DDX5, DDX3X, VIM, HSP90, and hnRNP F, was previously reported using a double hybrid strategy ^38, 39^ Additionally, HSP90, DDX5, and PRDX6, identified as NS5 interacting proteins, have been described to be associated with NS3, which in turn is also associated to NS5 during viral replication^39, 40^. Interestingly, the HSP90 protein was previously identified from our group as a putative DENV receptor in the monocytic cell line U937^41^. Another important group of proteins that we identified as NS5 interacting proteins were the hnRNPs. Some of them have been reported as important factors during DENV replicative cycle such as hnRNP C1/C2, hnRNP H, and hnRNP K^26, 27^, while others such as eEF2, DDX5 and DDX3X have been reported and characterized previously participating in the replicative cycle of other viruses^42^. Some cellular proteins bound to NS5 identified by De Maio et al., using nuclear extracts, although different from the ones identified here, belong to the same family. For example, they reported the hnRNP A0 and hnRNP R; the helicases DDX23 and DDX21; and the splicing factors SF3A2 and SF3B1, while we identified hnRNP F and hnRNP K; DDX5 and DDX3X and SFPQ. The main difference between both assays is that we used a recombinant protein and not the viral protein produced during infection.

Since DDX5 was previously identified as NS5 interacting protein and participates in the replicative cycle of other viruses^43–46^, we analyzed in further detail the importance and behavior of DDX5 protein during DENV infection. DDX5 belongs to the DEAD-box RNA helicase (Asp-Glu-Ala-Asp) family that plays crucial roles in many cellular processes^43^. Specifically, DDX5 is an important transcriptional factor and a co-activator during cell proliferation and differentiation^47^ and it is considered a multifunctional protein because it unwinds double-stranded RNA and is a nucleocytoplasmic shuttling protein whose action is mediated by Ran GTPase^48^.

Our results of immunoprecipitation and eletron microscopy indicate that DDX5 interacts with NS5 in the nucleus and cytoplasm of infected cells. Although it is not clear if the association between DDX5 and NS5 is direct or indirect, we know that it is not RNA mediated because cellular extracts used for the immunoprecipitation were treated with micrococcal endonuclease, eliminating the protein interactions mediated by RNA.

Although the specific role of DDX5 in the cytoplasm of DENV infected cells is unclear, our results indicate that it may be required for an IFN dependent response, since its relocation to the cytoplasm and degradation correlates with the evasion of the immune response. Similarly, DDX21, which activates the innate immune response, is also redistributed from the nucleus to the cytoplasm during infection with DENV and it is degraded by the action of the NS2B-NS3 protease^23^. This fact suggests that flavivirus such as DENV hijack some of the immune response targeting proteins present in the nucleus, and relocate them to the cytoplasm promoting its degradation. Interestingly, some other nuclear proteins such as DDX56, PTB, the autoantigen La hnRNP K, hnRNP C/C1, the high mobility box 1 protein (HMGB1), and TIA1/TIAR are also relocated from the nucleus to the cytoplasm during flavivirus infection^26, 31–34, 44–46, 49–52^. In all these cases, its translocation is necessary for an efficient viral replication or to inhibit the immune response.

Considering that the proteins that we identified as well as many others change their location from the nucleus to the cytoplasm during flavivirus infection and that viruses have the ability to hijack important components of the host cell, we analyzed the possible alteration in the function of the NPC, the main regulator of the nuclear-cytoplasmic transport. Our results indicate that during DENV infection the integrity and distribution of at least Nup153, Nup98, and Nup62 was disrupted, while during ZIKV infection the integrity of TPR, Nup153 and Nup98 was altered as well as the distribution of Nup62, being the viral serine protease NS2B-NS3 of both DENV and ZIKV the responsible molecules for these NUPs cleavage. This is the first report of the effect of flavivirus infection in this important nuclear structure. Some other cytoplasmic viruses utilize the NPC as an attractive anti-host target. Picornaviruses are particularly expert in disturbing the NPC; for example, the amino-terminal Leader protein (L) from the encephalomyocarditis virus (EMCV) induces hyperphosphorylation of Nup62, Nup153, and TPR, through a Ran-dependent, mitogen-activated protein kinase cascade^53^ disrupting their activity, while the viral protease 2Apro of poliovirus and rhinovirus cleaves Nup62, Nup98, and Nup153^54–57^ disrupting NPC activity. It is well known that the differences in NUPs cleavage cause differential inhibition of nuclear transport pathways through NPC^58, 59^. One of the nuclear transport pathways inhibited by the disruption of some FG-NUPs induced by DENV and ZIKV was the mRNAs export. The inhibition of this transport was first described in DENV infected cells as a consequence of splicing inhibition induced by NS5^14^; however, it is evident from our results and from the results obtained by De Maio et al, that the inhibition of poly A mRNAs export is a consequence of splicing inhibition induced by NS5 and the disruption in the NPC caused by NS2B-NS3, because the solely transfection of NS2B-NS3, which causes a disruption in FG-NUPs inhibited mRNA export dramatically, but not the transfection of the mutated protease, which does not alter FG-NUPs. Thus, as it happens with other processes, flavivirus hijack the main regulator of the nuclear-cytoplasmic transport which is the NPC inducing a drastic inhibition in the export of mRNA. This inhibition provides to flavivirus an advantage in the translation of its own genome because the ribosomes are more accessible to translate mainly viral RNAs. On the other hand, many signaling pathways end with the translocation of transcription factors to the nucleus to induce anti-viral gene expression; under infection conditions, even though the transcription factors could be transported to the nucleus and transcribe new genes, the possibility of being exported to be translated and induce the anti-viral conditions is reduced^60^.

Analyzing the time of infection in which NUPs degradation occurred, we found that Nup98 is the first NUP cleaved (from 12 h for ZIKV and at 24 h for DENV). Interestingly, it has been described that Nup98 is also a transcription factor involved in the expression of immune response genes^61, 62^ This would suggest that its degradation alters not only the mRNA export but also the expression of Nup98-dependent genes such as CDK9, RNAPII, and HLA-DRA, which in turn regulate the expression of TNF-alpha, inducible IL-6, and IFN-gamma^62^. All these cytokines are relevant during flavivirus infection^63^ and its abundance has to be evaluated after transfection with WT and mutant viral proteases.

In this work, several lines of evidence indicate that the viral protease NS2B-NS3 is responsible for NUPs cleavage. First, the serine protease inhibitors, TLCK and Leupeptin, prevented Nup98 and Nup62 cleavage. Second, the size of the predicted cleaved products from Nup98 and Nup62 correlated with those of the products obtained after DENV infection. Third, the transfection of DENV NS2B-NS3 protease was sufficient to inhibit the nuclear ring recognition detected in mock-infected cells with the Mab414 antibody. Fourth, the mutant but not the WT protease was unable to cleave NUPs in transfected cells. Interestingly, the viral protease is located early after infection (8 hrs for DENV and 12 hrs for ZIKV) in the perinuclear region as well as in the nucleus of the cells and at 24 hpi the protein is present in the cytoplasm in the perinuclear region. Degradation of nucleoporins by the activity of viral proteases located in the perinuclear region, such as the 2A protease from poliovirus, or in the nucleus such as the 3CD protease-polymerase from rhinovirus^35, 36^ have been reported, suggesting that NS3 located in this two particular locations could reach and cut its target NUPs.

Up to now, there was no evidence that NS3 could reach the nucleus of mammalian infected cells; however, it was observed in the nucleus in mosquito cells infected with DENV and ZIKV^64^. Considering the possible presence of NS3 in the nucleus, we analyzed if NS3 could contain a putative Nuclear Localization Sequence (NLS). Two types of NLSs have been described: the classical (cNLS) that can be mono and bipartite, and the non-classical (ncNLS). One important feature of the cNLS is the presence of basic residues of Arginine and Lysine (K and R)^65^.

Using two different Software: NLS-Mapper^19^ and Seq-NLS University South Carolina^20^, an NLS was predicted between amino acids 201 to 217 of the NS3 protein of the four DENV serotypes and ZIKV (Figure 6A Supplementary material). These sequences are located within the helicase domain. Additionally, using the LocNes software^21^ a Nuclear Export Sequence (NES) was predicted between amino acids 270 to 282 of the NS3 protein of the four DENV serotypes and ZIKV, also within the helicase domain (Figure 6A Supplementary material). Interestingly, the NES shows high conservation of hydrophobic residues (Leu, Ile, Val, and Met)^66^. The presence of both sequences within the NS3 protein of the four DENV serotypes and ZIKV suggests that NS3 is able to shuttle between the nucleus and the cytoplasm during infection. To confirm this suggestion, mutations in the NLS and NES of both recombinant proteases should be generated.

Although the integrity and distribution of FG-NUPs are required for an efficient nuclear-cytoplasmic transport, the partial or total degradation of some of them could be altering specific nuclear-cytoplasmic pathways. In this regard, even though our results clearly indicate that the export of mRNAs is blocked, the export pathway of DDX5 and hnRNP F is unaltered. This finding is particularly interesting because DDX5, hnRNP F, and NS5 proteins use the same pathway to move from the nucleus to the cytoplasm; thus, the import and export pathways used by NS5 guarantee its own and some other proteins movement between these two cellular compartments.

Our results showed a new function of the NS5 protein in the translocation from the nucleus to the cytoplasm of at least two proteins: DDX5 and hnRNP F. Although we only analyzed the participation of NS5 in the export of these two particular proteins, it is possible that NS5 is playing an active role in the shuttling between the nucleus and the cytoplasm of some other proteins during flavivirus infection such as PTB (data not shown).

Here we also found that the DDX5 is relocated to the cytoplasm by NS5 and degraded by NS3, in accordance with a previous report that showed that DDX21 protein, which activates the innate immune response, is redistributed from the nucleus to the cytoplasm and degraded by the direct or indirect action of the NS2B-NS3 protease during infection with DENV^23^. It is possible that the relocation to the cytoplasm of some nuclear proteins could play two different functions: 1) recruit cellular proteins required for a viral replicative cycle, and 2) highjack in the cytoplasm factors required for the expression of molecules involved in immune response, to control the antiviral response. Silencing of DDX5 showed that this protein is required for type I IFN signaling since in its absence, the cell was unable to induce the expression of MHC in cells treated with poly I:C. Recently the role of DDX5 in antiviral responses during HBV and Myxovirus infection has been reported^30^, however, more studies are required to understand the specific function of all these factors in flavivirus replicative cycle.

In summary, our results describe for the first time that the infection with flaviviruses such as DENV and ZIKV induces the disruption and degradation of some FG-NUPs by the activity of NS2B-NS3 protease, causing an inhibition in mRNA export and innate immune response signaling. Interestingly, the disruption of some of the FG-NUPs induced by DENV and ZIKV does not cause a massive mobilization of components between the nucleus and the cytoplasm, because while a dramatic inhibition in RNA export was confirmed, the export of some cellular proteins such as DDX5 and hnRNP F was not modified. A surprising result was the ability of NS5 to induce the relocation of DDX5 and hnRNP F to the cytoplasm, supporting the idea that NS3 and NS5 play together important functions in the nuclear-cytoplasmic transport; first, disrupting some FG-NUPs and altering some nuclear-cytoplasmic transport pathways and mediating export of some nuclear proteins respectively. It is likely that these two functions have a significant impact in the viral replicative cycle as well as in the modulation of the immune response.

Finally, we propose a hypothetical model of the role of NS5 and NS3 in infected cells. Early after infection NS5 is re-localized by alpha-beta importin systems into the nucleus and NS3 is located in nucleus and in the perinuclear region of the cell. NS3 degrades FG-NUPs altering mRNAs export and NS5 highjacks some nuclear proteins (DDX5, and hnRNP F), which are exported to the cytoplasm probable by the CRM1 pathway. In the cytoplasm NS3 degrades DDX5 inhibiting IFN response (Figure 14).

**Figure 14.**
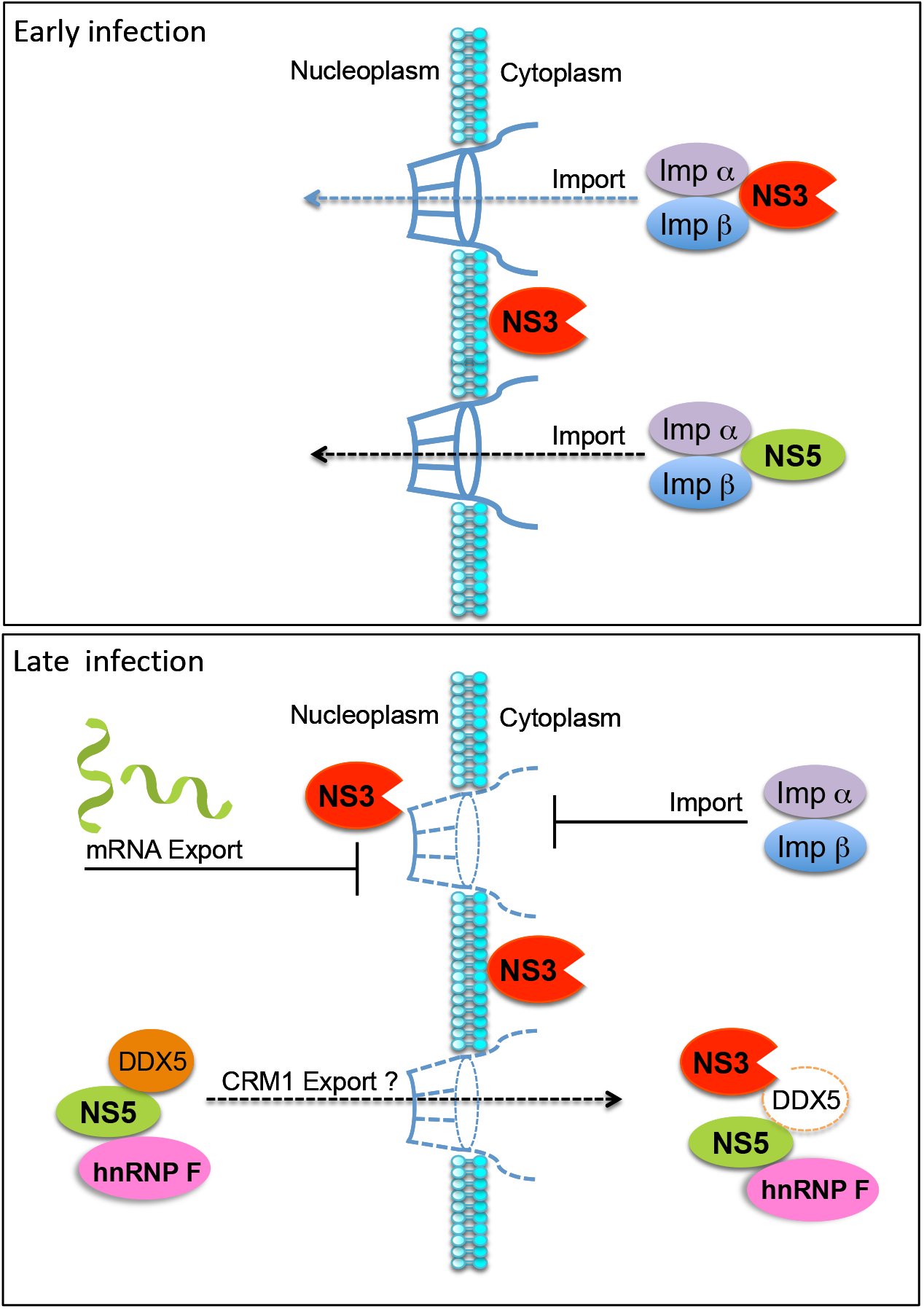
Hypothetical model of the role of NS5 and NS3 in infected cells. Early after infection NS3 and NS5 viral proteins are re-localized by alpha-beta importin into the nucleus. NS3 degrades FG-NUPs altering mRNAs export and NS5 highjacks some nuclear proteins such as DDX5 and hnRNP F, which are exported to the cytoplasm probable by CRM1 pathway. In the cytoplasm NS3 degrades DDX5 inhibiting IFN response.

| **identified proteins** | **Mascot score** | **# Peptide sequences identified** | **String Accession #** |
| --- | --- | --- | --- |
| Tubulin beta chain | 299 | 7 | TBB5 |
| Tubulin beta-2A chain | 191 | 6 | TBB2A |
| Tubulin beta-4B chain | 160 | 5 | TBB4B |
| Tubulin alpha-1A chain | 38 | 5 | TBA1A |
| Vimentin | 286 | 16 | VIME |
| L-lactate dehydrogenase B chain | 76 | 2 | LDHB |
| Fructose-bisphosphate aldolase A | 63 | 3 | ALDOA |
| Endoplasmin | 59 | 7 | ENPL |
| Probable ATP-dependent RNA helicase DDX5 | 59 | 2 | DDX5 |
| Heat shock protein HSP 90-beta | 54 | 5 | HS90B |
| Glyceraldehyde-3-phosphate dehydrogenase | 53 | 1 | G3P |
| Far upstream element-binding protein 2 | 52 | 1 | FUBP2 |
| Elongation factor 2 | 52 | 1 | EF2 |
| Insulin-like growth factor 2 mRNA-binding protein | 49 | 2 | IF2B2 |
| 60 kDa heat shock protein, mitochondrial | 45 | 3 | CH60 |
| ATP synthase subunit alpha, mitochondrial | 45 | 3 | ATPA |
| cAMP-dependent protein kinase type I-alpha regulatory subunit | 45 | 1 | KAP0 |
| Peroxiredoxin-1 | 43 | 4 | PRDX1 |
| Peroxiredoxin-6 | 28 | 1 | PRDX6 |
| 40S ribosomal protein S25 | 49 | 1 | RS25 |
| 60S ribosomal protein L18 | 41 | 1 | RL18 |
| 60S ribosomal protein L19 | 40 | 1 | RL19 |
| 40S ribosomal protein S14 | 56 | 1 | RS14 |
| 60S ribosomal protein L27 | 29 | 1 | RL27 |
| Prelamin-A/C OS | 39 | 4 | LMNA |
| Cofilin-1 | 39 | 3 | COF1 |
| Pyruvate kinase isozymes M1/M2 | 38 | 3 | KPYM |
| Heterogeneous nuclear ribonucleoprotein F | 36 | 1 | HNRPF |
| Heterogeneous nuclear ribonucleoprotein K | 36 | 4 | HNRPK |
| ATP-dependent RNA helicase DDX3X | 33 | 1 | DDX3X |
| Histone H2B type 1-C/E/F/G/I | 31 | 2 | H2B1C |
| Stress-70 protein, mitochondrial | 31 | 1 | GRP75 |
| Aconitate hydratase, mitochondrial | 30 | 1 | ACON |
| Heat shock protein beta-1 | 28 | 1 | HSPB1 |
| Splicing factor, proline- and glutamine-rich | 27 | 1 | SFPQ |
| 14-3-3 protein zeta/delta | 27 | 4 | 1433Z |

## Acknowledgments

The authors thank to Fernando Medina and Jaime Zarco for their technical assistance.

## REFERENCES

1. Han P, Ye W, Lv X, et al. DDX50 inhibits the replication of dengue virus 2 by upregulating IFN-beta production. Arch Virol 2017; 162(6): 1487–94.

2. Morrison J, Aguirre S, Fernandez-Sesma A. Innate immunity evasion by Dengue virus. Viruses 2012; 4(3): 397–413.

3. Soto-Acosta R, Bautista-Carbajal P, Cervantes-Salazar M, Angel-Ambrocio AH, del Angel RM. DENV up-regulates the HMG-CoA reductase activity through the impairment of AMPK phosphorylation: A potential antiviral target. PLoS pathogens 2017; 13(4).

4. Malet H, Masse N, Selisko B, et al. The flavivirus polymerase as a target for drug discovery. Antiviral research 2008; 80(1): 23–35.

5. Deubel V, Kinney RM, Trent DW. Nucleotide sequence and deduced amino acid sequence of the nonstructural proteins of dengue type 2 virus, Jamaica genotype: comparative analysis of the full-length genome. Virology 1988; 165(1): 234–44.

6. Dong H, Chang DC, Hua MH, et al. 2’-O methylation of internal adenosine by flavivirus NS5 methyltransferase. PLoS pathogens 2012; 8(4): e1002642.

7. Egloff MP, Benarroch D, Selisko B, Romette JL, Canard B. An RNA cap (nucleoside-2’-O-)-methyltransferase in the flavivirus RNA polymerase NS5: crystal structure and functional characterization. The EMBO journal 2002; 21(11): 2757–68.

8. Iglesias NG, Filomatori CV, Gamarnik AV. The F1 motif of dengue virus polymerase NS5 is involved in promoter-dependent RNA synthesis. Journal of virology 2011; 85(12): 5745–56.

9. Potisopon S, Priet S, Collet A, Decroly E, Canard B, Selisko B. The methyltransferase domain of dengue virus protein NS5 ensures efficient RNA synthesis initiation and elongation by the polymerase domain. Nucleic acids research 2016; 44(6): 2974.

10. Ashour J, Laurent-Rolle M, Shi PY, Garcia-Sastre A. NS5 of dengue virus mediates STAT2 binding and degradation. Journal of virology 2009; 83(11): 5408–18.

11. Morrison J, Laurent-Rolle M, Maestre AM, et al. Dengue virus co-opts UBR4 to degrade STAT2 and antagonize type I interferon signaling. PLoS pathogens 2013; 9(3): e1003265.

12. Medin CL, Fitzgerald KA, Rothman AL. Dengue virus nonstructural protein NS5 induces interleukin-8 transcription and secretion. Journal of virology 2005; 79(17): 11053–61.

13. Mazzon M, Jones M, Davidson A, Chain B, Jacobs M. Dengue virus NS5 inhibits interferon-alpha signaling by blocking signal transducer and activator of transcription 2 phosphorylation. The Journal of infectious diseases 2009; 200(8): 1261–70.

14. De Maio FA, Risso G, Iglesias NG, et al. The Dengue Virus NS5 Protein Intrudes in the Cellular Spliceosome and Modulates Splicing. PLoS pathogens 2016; 12(8): e1005841.

15. Hannemann H, Sung PY, Chiu HC, et al. Serotype-specific differences in dengue virus non-structural protein 5 nuclear localization. The Journal of biological chemistry 2013; 288(31): 22621–35.

16. Pryor MJ, Rawlinson SM, Butcher RE, et al. Nuclear localization of dengue virus nonstructural protein 5 through its importin alpha/beta-recognized nuclear localization sequences is integral to viral infection. Traffic 2007; 8(7): 795–807.

17. Tay MY, Smith K, Ng IH, et al. The C-terminal 18 Amino Acid Region of Dengue Virus NS5 Regulates its Subcellular Localization and Contains a Conserved Arginine Residue Essential for Infectious Virus Production. PLoS pathogens 2016; 12(9): e1005886.

18. Anglero-Rodriguez YI, Pantoja P, Sariol CA. Dengue virus subverts the interferon induction pathway via NS2B/3 protease-IkappaB kinase epsilon interaction. Clinical and vaccine immunology : CVI 2014; 21(1): 29–38.

19. Kosugi S, Hasebe M, Tomita M, Yanagawa H. Systematic identification of cell cycle-dependent yeast nucleocytoplasmic shuttling proteins by prediction of composite motifs. Proceedings of the National Academy of Sciences of the United States of America 2009; 106(25): 10171–6.

20. Lin JR, Mondal AM, Liu R, Hu J. Minimalist ensemble algorithms for genome-wide protein localization prediction. BMC bioinformatics 2012; 13: 157.

21. Xu D, Marquis K, Pei J, et al. LocNES: a computational tool for locating classical NESs in CRM1 cargo proteins. Bioinformatics 2015; 31(9): 1357–65.

22. Dardenne E, Polay Espinoza M, Fattet L, et al. RNA helicases DDX5 and DDX17 dynamically orchestrate transcription, miRNA, and splicing programs in cell differentiation. Cell reports 2014; 7(6): 1900–13.

23. Dong Y, Ye W, Yang J, et al. DDX21 translocates from nucleus to cytoplasm and stimulates the innate immune response due to dengue virus infection. Biochem Biophys Res Commun 2016; 473(2): 648–53.

24. Li G, Feng T, Pan W, Shi X, Dai J. DEAD-box RNA helicase DDX3X inhibits DENV replication via regulating type one interferon pathway. Biochem Biophys Res Commun 2015; 456(1): 327–32.

25. Kumar R, Singh N, Abdin MZ, Patel AH, Medigeshi GR. Dengue Virus Capsid Interacts with DDX3X-A Potential Mechanism for Suppression of Antiviral Functions in Dengue Infection. Frontiers in cellular and infection microbiology 2017; 7: 542.

26. Brunetti JE, Scolaro LA, Castilla V. The heterogeneous nuclear ribonucleoprotein K (hnRNP K) is a host factor required for dengue virus and Junin virus multiplication. Virus Res 2015; 203: 84–91.

27. Mishra KP, Shweta, Diwaker D, Ganju L. Dengue virus infection induces upregulation of hn RNP-H and PDIA3 for its multiplication in the host cell. Virus Res 2012; 163(2): 573–9.

28. Noisakran S, Sengsai S, Thongboonkerd V, et al. Identification of human hnRNP C1/C2 as a dengue virus NS1-interacting protein. Biochem Biophys Res Commun 2008; 372(1): 67–72.

29. Tay MY, Fraser JE, Chan WK, et al. Nuclear localization of dengue virus (DENV) 1-4 non-structural protein 5; protection against all 4 DENV serotypes by the inhibitor Ivermectin. Antiviral research 2013; 99(3): 301–6.

30. Cheng W, Chen G, Jia H, He X, Jing Z. DDX5 RNA Helicases: Emerging Roles in Viral Infection. Int J Mol Sci 2018; 19(4).

31. Agis-Juarez RA, Galvan I, Medina F, et al. Polypyrimidine tract-binding protein is relocated to the cytoplasm and is required during dengue virus infection in Vero cells. The Journal of general virology 2009; 90(Pt 12): 2893–901.

32. Anwar A, Leong KM, Ng ML, Chu JJ, Garcia-Blanco MA. The polypyrimidine tract-binding protein is required for efficient dengue virus propagation and associates with the viral replication machinery. The Journal of biological chemistry 2009; 284(25): 17021–9.

33. Yocupicio-Monroy M, Padmanabhan R, Medina F, del Angel RM. Mosquito La protein binds to the 3’ untranslated region of the positive and negative polarity dengue virus RNAs and relocates to the cytoplasm of infected cells. Virology 2007; 357(1): 29–40.

34. Kamau E, Takhampunya R, Li T, et al. Dengue virus infection promotes translocation of high mobility group box 1 protein from the nucleus to the cytosol in dendritic cells, upregulates cytokine production and modulates virus replication. J Gen Virol 2009; 90(Pt 8): 1827–35.

35. Walker E, Jensen L, Croft S, et al. Rhinovirus 16 2A Protease Affects Nuclear Localization of 3CD during Infection. J Virol 2016; 90(24): 11032–42.

36. Amineva SP, Aminev AG, Palmenberg AC, Gern JE. Rhinovirus 3C protease precursors 3CD and 3CD’ localize to the nuclei of infected cells. J Gen Virol 2004; 85(Pt 10): 2969–79.

37. Ren Y, Seo HS, Blobel G, Hoelz A. Structural and functional analysis of the interaction between the nucleoporin Nup98 and the mRNA export factor Rae1. Proceedings of the National Academy of Sciences of the United States of America 2010; 107(23): 10406–11.

38. Khadka S, Vangeloff AD, Zhang C, et al. A physical interaction network of dengue virus and human proteins. Molecular & cellular proteomics : MCP 2011; 10(12): M111 012187.

39. Le Breton M, Meyniel-Schicklin L, Deloire A, et al. Flavivirus NS3 and NS5 proteins interaction network: a high-throughput yeast two-hybrid screen. BMC microbiology 2011; 11: 234.

40. Mairiang D, Zhang H, Sodja A, et al. Identification of new protein interactions between dengue fever virus and its hosts, human and mosquito. PloS one 2013; 8(1): e53535.

41. Reyes-Del Valle J, Chavez-Salinas S, Medina F, Del Angel RM. Heat shock protein 90 and heat shock protein 70 are components of dengue virus receptor complex in human cells. J Virol 2005; 79(8): 4557–67.

42. Owsianka AM, Patel AH. Hepatitis C virus core protein interacts with a human DEAD box protein DDX3. Virology 1999; 257(2): 330–40.

43. Kokolo M, Bach-Elias M. Downregulation of p68 RNA Helicase (DDX5) Activates a Survival Pathway Involving mTOR and MDM2 Signals. Folia biologica 2017; 63(2): 52–9.

44. Yocupicio-Monroy RM, Medina F, Reyes-del Valle J, del Angel RM. Cellular proteins from human monocytes bind to dengue 4 virus minus-strand 3’ untranslated region RNA. J Virol 2003; 77(5): 3067–76.

45. Garcia-Montalvo BM, Medina F, del Angel RM. La protein binds to NS5 and NS3 and to the 5’ and 3’ ends of Dengue 4 virus RNA. Virus Res 2004; 102(2): 141–50.

46. Chang CJ, Luh HW, Wang SH, Lin HJ, Lee SC, Hu ST. The heterogeneous nuclear ribonucleoprotein K (hnRNP K) interacts with dengue virus core protein. DNA Cell Biol 2001; 20(9): 569–77.

47. Ramanathan N, Lim N, Stewart CL. DDX5/p68 RNA helicase expression is essential for initiating adipogenesis. Lipids in health and disease 2015; 14: 160.

48. Zhou X, Luo J, Mills L, et al. DDX5 facilitates HIV-1 replication as a cellular cofactor of Rev. PloS one 2013; 8(5): e65040.

49. Dechtawewat T, Songprakhon P, Limjindaporn T, et al. Role of human heterogeneous nuclear ribonucleoprotein C1/C2 in dengue virus replication. Virology journal 2015; 12(1): 14.

50. Lee JH, Kim SH, Pascua PN, et al. Direct interaction of cellular hnRNP-F and NS1 of influenza A virus accelerates viral replication by modulation of viral transcriptional activity and host gene expression. Virology 2010; 397(1): 89–99.

51. Li W, Li Y, Kedersha N, et al. Cell proteins TIA-1 and TIAR interact with the 3’ stem-loop of the West Nile virus complementary minus-strand RNA and facilitate virus replication. J Virol 2002; 76(23): 11989–2000.

52. Reid CR, Hobman TC. The nucleolar helicase DDX56 redistributes to West Nile virus assembly sites. Virology 2017; 500: 169–77.

53. Porter FW, Brown B, Palmenberg AC. Nucleoporin phosphorylation triggered by the encephalomyocarditis virus leader protein is mediated by mitogen-activated protein kinases. Journal of virology 2010; 84(24): 12538–48.

54. Park N, Katikaneni P, Skern T, Gustin KE. Differential targeting of nuclear pore complex proteins in poliovirus-infected cells. Journal of virology 2008; 82(4): 1647–55.

55. Gustin KE, Sarnow P. Inhibition of nuclear import and alteration of nuclear pore complex composition by rhinovirus. Journal of virology 2002; 76(17): 8787–96.

56. Park N, Skern T, Gustin KE. Specific cleavage of the nuclear pore complex protein Nup62 by a viral protease. The Journal of biological chemistry 2010; 285(37): 28796–805.

57. Watters K, Inankur B, Gardiner JC, et al. Differential Disruption of Nucleocytoplasmic Trafficking Pathways by Rhinovirus 2A Proteases. Journal of virology 2017.

58. Wu X, Kasper LH, Mantcheva RT, Mantchev GT, Springett MJ, van Deursen JM. Disruption of the FG nucleoporin NUP98 causes selective changes in nuclear pore complex stoichiometry and function. Proc Natl Acad Sci U S A 2001; 98(6): 3191–6.

59. Tran EJ, Wente SR. Dynamic nuclear pore complexes: life on the edge. Cell 2006; 125(6): 1041–53.

60. Petersen JM, Her LS, Varvel V, Lund E, Dahlberg JE. The matrix protein of vesicular stomatitis virus inhibits nucleocytoplasmic transport when it is in the nucleus and associated with nuclear pore complexes. Molecular and cellular biology 2000; 20(22): 8590–601.

61. Panda D, Pascual-Garcia P, Dunagin M, et al. Nup98 promotes antiviral gene expression to restrict RNA viral infection in Drosophila. Proceedings of the National Academy of Sciences of the United States of America 2014; 111(37): E3890–9.

62. Panda D, Gold B, Tartell MA, Rausch K, Casas-Tinto S, Cherry S. The transcription factor FoxK participates with Nup98 to regulate antiviral gene expression. mBio 2015; 6(2).

63. Sam SS, Teoh BT, Chinna K, AbuBakar S. High producing tumor necrosis factor alpha gene alleles in protection against severe manifestations of dengue. International journal of medical sciences 2015; 12(2): 177–86.

64. Reyes-Ruiz JM, Osuna-Ramos JF, Cervantes-Salazar M, et al. Strand-like structures and the nonstructural proteins 5, 3 and 1 are present in the nucleus of mosquito cells infected with dengue virus. Virology 2018; 515: 74–80.

65. Liu P, Chen S, Wang M, Cheng A. The role of nuclear localization signal in parvovirus life cycle. Virology journal 2017; 14(1): 80.

66. la Cour T, Gupta R, Rapacki K, Skriver K, Poulsen FM, Brunak S. NESbase version 1.0: a database of nuclear export signals. Nucleic acids research 2003; 31(1): 393–6.

